# Excitatory drive to the globus pallidus external segment facilitates action initiation in non-human primates

**DOI:** 10.1101/2025.04.16.649214

**Authors:** Atsushi Yoshida, Okihide Hikosaka

**Author notes:** Correspondence: Atsushi Yoshida.

## Abstract

The external segment of the globus pallidus (GPe) has conventionally been regarded as a key relay in the indirect pathway of the basal ganglia, primarily mediating movement suppression; however, recent studies in rodents suggest a more complex role, including active facilitation of actions. Therefore, we investigated whether the primate GPe exhibits similar functional diversity by recording single-unit activity in two macaque monkeys performing a sequential choice task. This task separated processes of action initiation and suppression by requiring the monkeys to either accept a “good” object for reward or reject a “bad” object using one of multiple strategies. We identified three distinct neuronal clusters based on their firing patterns. Clusters 1 and 2 displayed elevated activity preceding contralateral saccades toward good objects, strongly correlating with shorter reaction times, suggesting a facilitative contribution. In contrast, Clusters 2 and 3 showed decreased activity during rejection of bad objects, reflecting proactive inhibition. Local pharmacological blockade of glutamate receptors within the caudodorsal GPe prolonged saccade latencies and reduced the frequency of rejection saccades, suggesting a causal role for excitatory drive in saccade facilitation. These findings expand the traditional view of the GPe beyond a purely inhibitory station, indicating that in primates it may simultaneously mediate both motor facilitation and proactive suppression. Our results emphasize the importance of characterizing circuit-specific and cell-type-specific roles of the GPe within basal ganglia networks, with implications for normal motor function and movement disorder pathophysiology under complex reward-based decision processes in non-human primates.

## Introduction

The basal ganglia are subcortical nuclei crucial for motor control, learning, and decision-making (1–3). Within these nuclei, the external globus pallidus (GPe) has long been viewed as a key element in the indirect pathway for motor suppression (2–5). According to this classical perspective, GPe neurons exert inhibitory gamma-aminobutyric acid (GABA)ergic influence over the subthalamic nucleus (STN), which in turn modulates thalamo-cortical circuits and affects the superior colliculus, a critical structure for oculomotor control.

However, recent cell-type-specific investigations—primarily in rodents— have challenged the notion that the GPe is simply a relay that mediates movement suppression (6–8). These studies highlight the GPe’s intricate cellular and circuit organization, revealing multiple neuronal populations (e.g., prototypical and arkypallidal neurons) with distinct molecular signatures and projection targets (9–12). Moreover, the GPe receives not only striatal and STN inputs, but also direct projections from the cortex and thalamus, calling into question the traditional view of a strictly unidirectional indirect pathway (13–17).

In contrast to these advances in rodent research, our understanding of GPe functions in non-human primates remains remarkably limited. Although advanced techniques for cell-type-specific manipulation have been slow to reach primate studies, certain electrophysiological findings suggest that the GPe cannot be characterized solely by its suppressive influence (18–20, 5). Recordings from GPe neurons in macaques reveal neuronal populations that exhibit increased activity during movement, suggesting a more complex role than simple suppression (21–23). For example, our own recordings in macaques revealed increased GPe activity during visually guided saccades and anti-saccades, pointing to a possible facilitative role in voluntary movements (24, 25). However, the anti-saccade task inherently involved both generating and suppressing responses, making it difficult to isolate the specific contribution of heightened GPe activity to movement facilitation.

Therefore, a critical question remains: “does the primate GPe actively promote voluntary movements via excitatory mechanisms, or does it primarily operate by suppressing or relaying inhibitory signals?” While rodent work has offered substantial clues about GPe heterogeneity and potential excitatory influences, conclusive data from non-human primates is lacking. To address this gap in the literature, we developed a novel sequential choice task specifically designed to disentangle facilitation from suppression by temporally separating these processes. In addition to recording time-locked GPe activity during key behavioral events, we employed local pharmacological manipulations to test whether excitatory drives to the GPe play a causal role in initiating movement. Specifically, we tested the hypothesis that, beyond its well-established function in suppressing unwanted actions, the primate GPe actively facilitates voluntary movements by integrating excitatory inputs, likely from the STN or cortex or both. We predicted that blocking these excitatory inputs would impair the initiation of voluntary movements. By taking this approach, we aimed to provide new insight into the dual role of the GPe in both inhibiting and facilitating voluntary movements, thus expanding the current understanding of basal ganglia function in primates.

## Results

### Behavioral performance in the sequential choice task

In this section, we demonstrate that monkeys effectively used learned object values to make accept or reject decisions, establishing a behavioral foundation for studying the GPe’s role in action selection.

We first confirmed that monkeys effectively utilized learned object values to accept or reject items in the sequential choice task, thereby establishing a robust behavioral basis for examining the GPe’s role in action selection. Two monkeys (Cr and Sp) were trained to perform a task (Figure 1A), in which they decided whether to accept a “good” (rewarding) object by making a saccade and holding fixation, or to reject a “bad” (non-rewarding) object using one of three strategies: “return” by briefly looking at the bad object and then returning to center, “stay” by maintaining central fixation, or “other” by making a saccade away from both the object and the center (Figures 1B and 1C).

**Figure 1.**
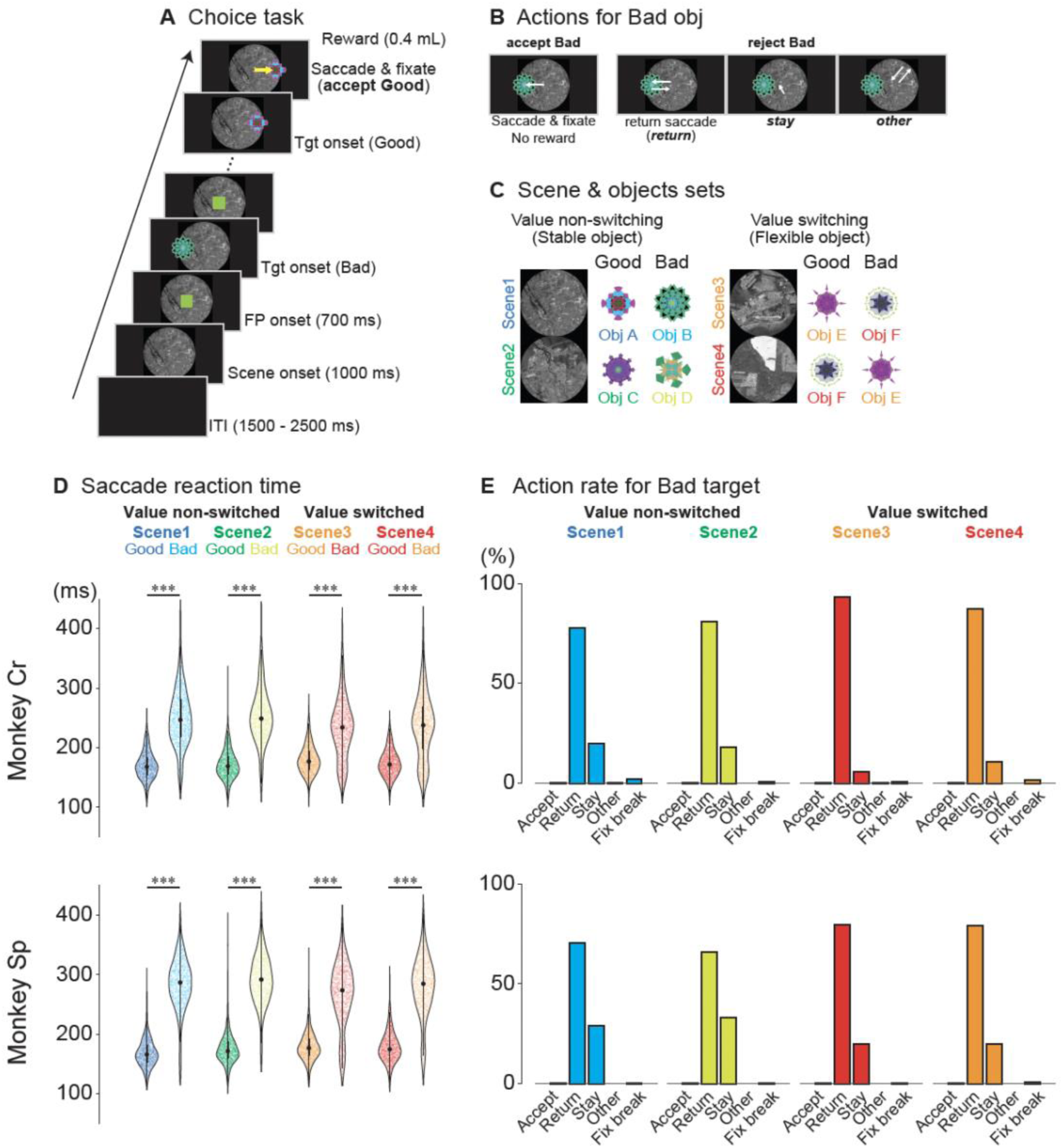
Choice task paradigm and behavioral performance of two macaque monkeys. (A) Schematic of the choice task. A background scene was initially presented (1000 ms), followed by a fixation point (FP, 700 ms). Subsequently, either a good (rewarded) or bad (non-rewarded) object was randomly presented, one at a time, at one of six possible locations (0°, 45°, 135°, 180°, 225°, or 315°). (B) Possible behavioral responses when a bad object was presented. When a bad object appeared, monkeys could either incorrectly accept it by making a saccade to the object and maintaining fixation on it (accept bad), or correctly reject it using one of three strategies: making a saccade toward the object and quickly returning to the central fixation point (return), maintaining fixation on the central fixation point without a saccade to the object (stay), or making a saccade in a direction away from both the object’s location and the central fixation point (other). (C) Scene and object combinations. Each recording session utilized one of six sets, with each set containing four scenes (see Figure S1 for all sets). Each scene was associated with two objects: one good (rewarded) and one bad (non-rewarded). In scenes 1 and 2, object values remained constant (value non-switching), while in scenes 3 and 4, the same objects were used but with reversed reward values (value-switching), illustrating context-dependent value assignment. (D) Saccadic reaction times for good and bad objects for the two monkeys (Cr and Sp). Violin plots show that, across all scenes and for both monkeys, saccades to good objects had significantly shorter latencies compared with saccades to bad objects during the return response (***p < 0.0001 for all comparisons, Welch’s t-tests). Each dot represents a single session. The thick horizontal line indicates the median, the box indicates the interquartile range (IQR), and the whiskers extend to the most extreme data point within 1.5 times the IQR. (E) Proportions of different behavioral responses to bad objects for each scene. Both monkeys exhibited similar patterns: incorrect acceptance of bad objects was rare, and rejection responses consisted predominantly of return saccades, followed by stay responses and then other responses. Accept, incorrect acceptance of bad objects; Return, return saccades; Stay, maintaining central fixation; Other, saccades away from both the object and the central fixation point; Fix break, fixation break errors. Abbreviations: FP, fixation point; ITI, inter-trial interval; Obj, object; Tgt, target.

As expected, the saccadic reaction times (RTs) were significantly shorter for good objects than for bad objects (Welch’s t-test, p < 0.0001; Table S1, Figure 1D), indicating that the monkeys successfully learned object values and used them to guide behavior. When rejecting bad objects, they primarily employed the “return” strategy (Figure 1E), accounting for most of the rejections (monkey Cr: 77.8– 93.4%; monkey Sp: 65.9–79.5%; Table S2), whereas “stay” was used less frequently (monkey Cr: 5.5–19.7%; monkey Sp: 20.0–33.2%). This preference for the “return” strategy likely reflects task timing, since “stay” required 400 ms of fixation, whereas “return” allowed for immediate presentation of the next target and potentially faster trial progression. We also observed that the proportion of “stay” responses differed between stable-value scenes (1 and 2) and flexible, context-dependent scenes (3 and 4), with significantly more “stay” responses in the stable-value scenes (Fisher’s exact test: monkey Cr, p = 9.76×10^−51^; monkey Sp, p = 1.66×10^−28^). This pattern suggests that higher cognitive demands for processing context-dependent values in scenes 3 and 4 reduced the tendency to remain fixated.

Together, these findings confirmed that the monkeys discriminated between good and bad objects and strategically chose between multiple rejection options, laying the groundwork for our subsequent investigation of the GPe’s contribution to action selection.

### General characteristics of GPe neuronal activity during the choice task

To investigate the role of the GPe neurons in action selection, we analyzed their firing rates and temporal dynamics during key task events: scene onset, target onset, and saccade onset. This analysis of overall GPe activity provided a basis for identifying distinct functional subpopulations in subsequent sections.

We recorded single-unit activity from 205 task-responsive GPe neurons (Monkey Cr: 111; Monkey Sp: 94) during the sequential choice task, focusing on trials with contralateral object presentations (see Methods for details). When we aligned neuronal activity to three key task events—scene, target, and saccade onset—we found that many GPe neurons displayed distinct firing modulations at one or more of these time points. Some neurons exhibited an increase in firing immediately after scene onset, whereas others showed changes time-locked to target onset or saccade initiation.

Taken together, these preliminary observations suggest the existence of multiple functional subpopulations within the GPe, prompting us to examine whether such diverse activity patterns reflect distinct roles in movement facilitation and inhibition.

### Classification of GPe neurons into three clusters based on firing patterns during the choice task

In this section, we performed k-means clustering to identify functional subpopulations of GPe neurons and lay the foundation for analyzing their roles in movement facilitation and inhibition.

To further characterize the functional heterogeneity of GPe neurons, we used k-means clustering to classify neurons based on their average standardized firing rates during the presentation of contralateral good and bad objects (100–300 ms after object onset). We chose k-means clustering as it provides an objective, data-driven, and comprehensive approach to classifying neurons. This method minimizes researcher bias by grouping neurons based on the similarity of their firing patterns, without pre-defined categories or assumptions about the underlying functional organization of the GPe. Furthermore, k-means clustering ensures that all recorded neurons are assigned to a cluster, providing a comprehensive classification of the recorded population. This analysis revealed three distinct clusters (Figure S2A and S2B): Cluster 1 (n = 58), Cluster 2 (n = 86), and Cluster 3 (n = 61). Briefly, Cluster 1 neurons showed increased activity after both scene and target onset, encoding both object value and spatial location. Cluster 2 neurons also exhibited increased activity after scene onset, but showed distinct firing patterns at target onset, increasing firing for good objects and decreasing for bad objects. Cluster 3 neurons showed little change in activity at scene onset, but decreased firing at target onset, particularly for bad objects. Furthermore, these clusters exhibited different baseline firing rates, with Cluster 1 showing a significantly lower rate than Clusters 2 and 3.

### Comparison of electrophysiological properties among the three GPe neuronal clusters

Herein, we compare the spike waveforms and baseline firing rates across the three clusters to determine whether each subgroup exhibited unique electrophysiological signatures.

Having identified three functionally distinct clusters, we subsequently asked whether these subpopulations exhibited unique electrophysiological signatures, such as differences in spike waveforms or baseline firing rates. To address this, we focused on spike waveform duration and baseline firing rate. While subtle but significant discrepancies in spike waveform duration emerged between Cluster 1 and 2 (Kruskal–Wallis test, H = 8.98, p = 1.01 × 10^−2^; Dunn’s test, p = 8.52 × 10^−3^), the overall spike shapes across all three clusters remained broadly similar (Figures S2C and S2D).

We also compared baseline firing rates across clusters and observed a significant overall difference (Kruskal–Wallis test, H = 14.82, p = 6.05 × 10^−5^; Figure S2E). Post-hoc analyses (Dunn’s test) revealed that Cluster 1 had a markedly lower baseline firing rate compared with both Cluster 2 and 3, whose rates did not differ from each other. The mean firing rates were 42.9 ± 22.4 Hz, 59.2 ± 27.7 Hz, and 58.4 ± 29.3 Hz for clusters 1, 2, and 3, respectively.

Together, these electrophysiological distinctions suggest that each cluster may represent a functionally distinct subpopulation within the GPe, potentially reflecting different cellular subtypes or projection targets.

### Modulation of neuronal activity in the three GPe clusters by scene and target onset

In this section, we examined how each cluster responded to scene and target onset, focusing on differences in value and spatial coding.

To clarify how each cluster encodes both the visual context (scene onset) and the specific target (object onset), we aligned GPe neuron activity to these two critical time points and examined how firing differed by object value and location. We focused on the normalized population activity across clusters (Figure 2, Tables S3-S5).

**Figure 2.**
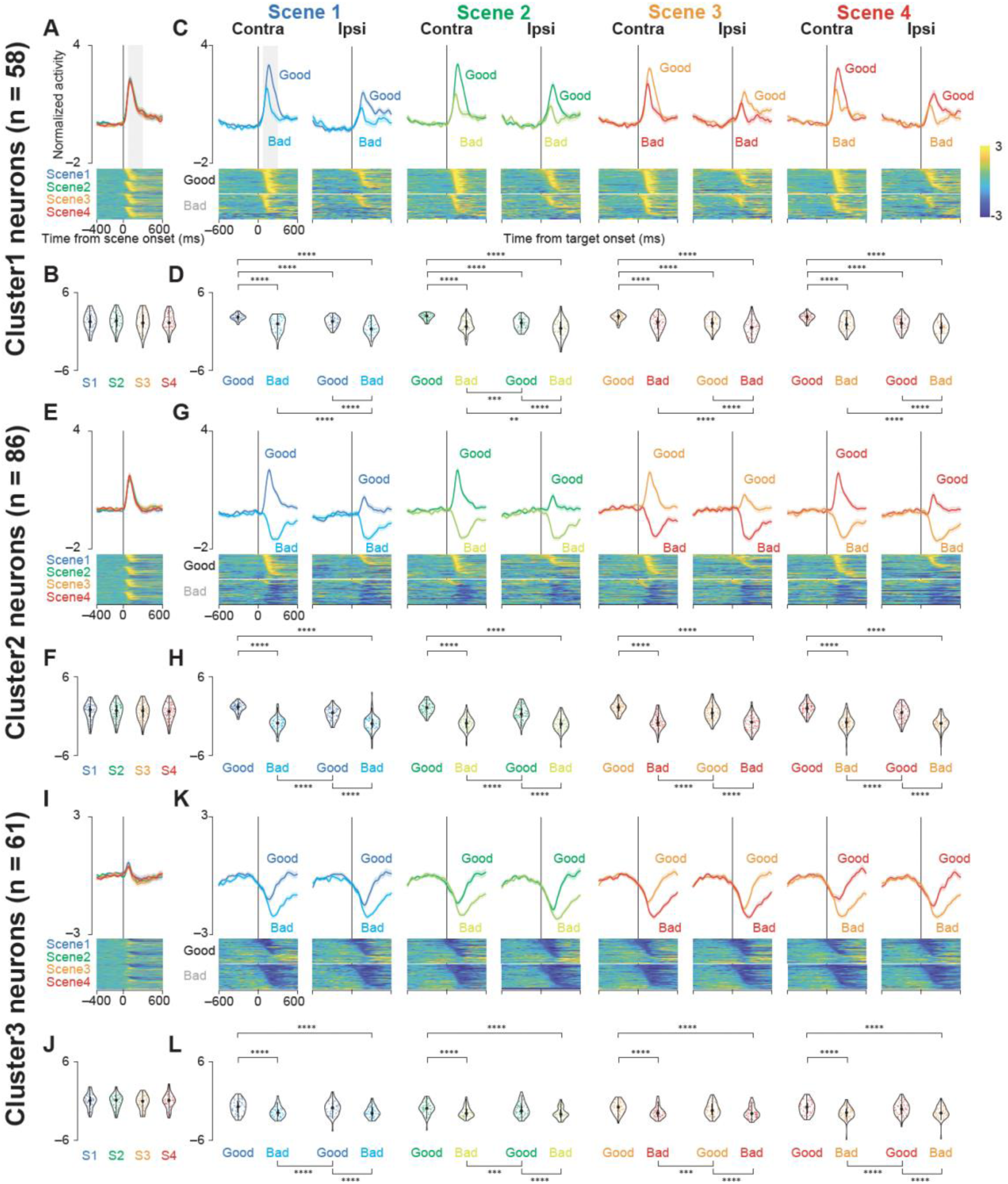
Population activity of three groups of GPe neurons at scene and target onsets during the choice task. (A, E, I) Mean normalized population activity (activity was normalized by subtracting the baseline firing rate and dividing by the standard deviation) of Cluster 1 (A), 2 (E), and 3 (I) neurons aligned to the onset of the scene (scenes 1–4) during the choice task. The shaded areas indicate ± SEMs. The lower panels show color maps of the normalized activity of individual neurons. Each row in the color maps represents a single neuron, and the neurons are sorted based on the time at which their standardized activity exceeded a threshold (see Methods for details). (B, F, J) Violin plots showing the distribution of the mean normalized neuronal activity of individual neurons in Clusters 1 (B), 2 (F), and 3 (J) for each scene onset during the choice task. Neuronal activity was measured for a 200-ms interval beginning 100 ms after scene onset (gray rectangle in A, E, and I). The larger circle indicates the median value, the thick vertical line shows the interquartile range (IQR), and the thin vertical line indicates the range from the lower to the upper adjacent values (1.5xIQR below the first quartile and 1.5xIQR above the third quartile, respectively). (C, G, K) Mean normalized population activity of Cluster 1 (C), 2 (G), and 3 (K) neurons aligned to the onset of the contralateral and ipsilateral good and bad targets for scenes 1– 4 during the choice task. The shaded areas indicate ± SEMs. The lower panels show color maps of the normalized neuronal activity of individual neurons, with the neurons sorted in the same order as in A, E, and I. (D, H, L) Violin plots showing the distribution of the mean normalized neuronal activity of individual neurons in Clusters 1 (D), 2 (H), and 3 (L) for each target onset condition during the choice task. Neuronal activity was measured for a 200-ms interval beginning 100 ms after target onset (gray rectangle in C, G, and K). The format of the violin plots is the same as in B, F, and J. The asterisks indicate significant differences in neuronal activity among the four conditions (contra good, contra bad, ipsi good, ipsi bad) within each scene (post-hoc pairwise t-tests with Bonferroni correction, *p < 0.05, **p < 0.01, ***p < 0.001, ****p < 0.0001). The details of the statistical tests are summarized in Tables S3-S5. Abbreviations: GPe, external segment of the globus pallidus; SEM, standard errors of the mean.

Cluster 1 showed increased firing following scene onset; however, this elevation did not differ significantly among the four scenes (Figures 2A and 2B; parametric bootstrap test, p = 0.51). In contrast, when aligned to target onset, Cluster 1 neurons displayed significantly different activity across the four conditions (contralateral good/bad, ipsilateral good/bad) in every scene (Figures 2C and 2D; p < 0.001; Table S3). Specifically, they increased firing for both good and bad objects (with a stronger response to good objects), and showed a preference for contralateral presentations. These patterns suggest that Cluster 1 neurons encode both object value and spatial location.

Cluster 2 behaved similarly to Cluster 1 at scene onset, showing no difference among the four scenes (Figures 2E and 2F; p = 0.78). However, on target onset, Cluster 2 firing differed markedly across conditions in all scenes (Figures 2G and 2H; p < 0.001; Table S4). Unlike Cluster 1, Cluster 2 firing increased for good objects but decreased for bad objects, responding more strongly to contralateral good targets while showing no strong laterality preference for bad targets. This pattern underscores a prominent role for value processing in Cluster 2.

Cluster 3 neuron activity at scene onset remained unchanged among the four scenes (Figures 2I and 2J; p = 0.35). In contrast, when aligned to target onset, Cluster 3 firing differed significantly for contralateral and ipsilateral good or bad objects (Figures 2K and 2L; p < 0.001; Table S5). In this cluster, firing generally decreased for both good and bad objects, with a more pronounced suppression for bad objects. As scene-onset activity remained relatively constant across the four scenes, it appears that Cluster 3 modulation is driven primarily by object value, rather than general visual factors.

Overall, these results indicate that each cluster exhibits a distinct combination of spatial and value coding, illuminating multiple functional pathways through which the GPe might integrate context and object significance during decision-making.

### Activity of the three GPe neuronal clusters aligned to saccade onset

Next, we investigated the relationship between neuronal firing and saccade initiation, highlighting each Cluster’s potential role in movement execution.

To determine whether neuronal firing in the three clusters related to saccade initiation, we aligned their activity to saccade onset and compared responses for contralateral and ipsilateral good or bad objects. This analysis revealed distinct firing patterns across clusters (Figure 3, Tables S6–S8).

**Figure 3.**
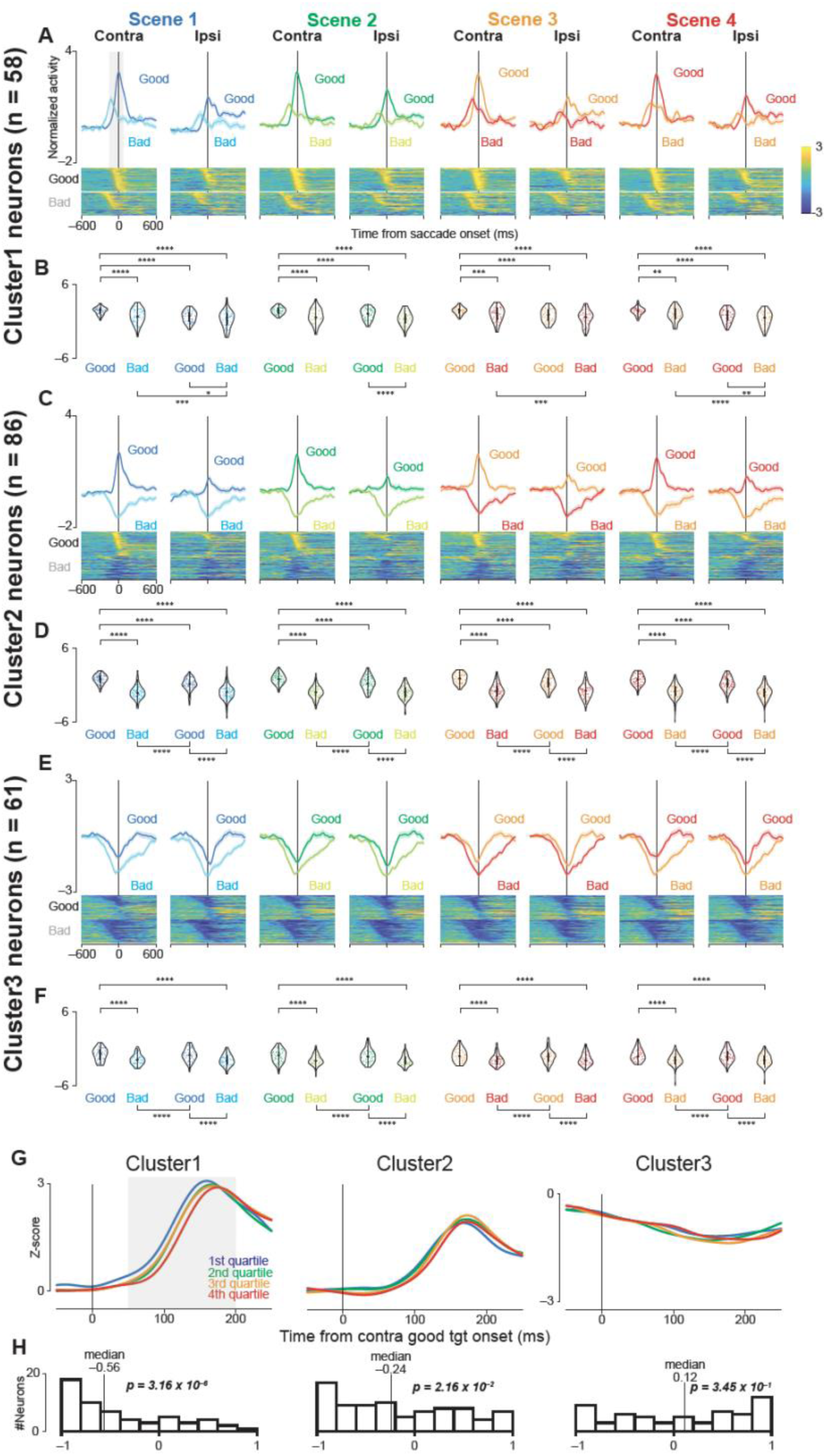
Population activity of three groups of GPe neurons aligned to saccade onset during the choice task. (A, C, E) Mean normalized population activity (activity was normalized by subtracting the baseline firing rate and dividing by the standard deviation) of Cluster 1 (A), 2 (C), and 3 (E) neurons aligned to saccade onset for contralateral or ipsilateral, good or bad objects in all scenes during the choice task. The shaded areas indicate ± SEMs. The lower panels show color maps of the normalized neuronal activity of individual neurons, with each row representing a single neuron. The neurons are sorted in the same order as in Figures 2A, 2E, and 2I. (B, D, F) Violin plots showing the distribution of the mean normalized neuronal activity of individual neurons in Clusters 1 (B), 2 (D), and 3 (F) when the monkeys made a saccade to the target during the choice task. Neuronal activity was measured for a 200-ms interval from 150 ms before to 50 ms after saccade onset (gray rectangle in A, C, and E). The format of the violin plots is the same as in Figures 2B, 2F, and 2J. The asterisks indicate significant differences in neuronal activity among the four conditions (contra good, contra bad, ipsi good, ipsi bad) within each scene (post-hoc pairwise t-tests with Bonferroni correction, *p < 0.05, **p < 0.01, ***p < 0.001, ****p < 0.0001). The details of the statistical tests are summarized in Tables S6-S8. (G) Time courses of the normalized population activity of neurons in Clusters 1, 2, and 3 when contralateral good objects were presented. For individual neurons, the trials were divided into four groups based on the saccade reaction times: 1st quartile (fastest, blue), 2nd quartile (green), 3rd quartile (orange), and 4th quartile (slowest, red). Each line represents the average activity of all neurons in each cluster for each quartile group. (H) Histograms showing the distribution of correlation coefficients between neuronal activity and saccade reaction times for each neuron in clusters 1, 2, and 3. Neuronal activity was measured for a 150-ms interval beginning 50 ms after target onset (gray rectangle in G). The vertical lines indicate the median values. The p-values are from Wilcoxon signed-rank tests, testing whether the median correlation coefficient is significantly different from zero. Abbreviations: GPe, external segment of the globus pallidus; SEM, standard errors of the mean.

Cluster 1 activity differed significantly among all four conditions (contralateral good/bad, ipsilateral good/bad) in every scene (Figures 3A and 3B; parametric bootstrap test, p < 0.001; Table S6). Overall, firing was higher for saccades to good objects than bad objects, and higher for contralateral rather than ipsilateral directions. Such a consistent elevation of firing suggests that Cluster 1 contributes to saccade generation.

Cluster 2 also showed significantly different activity across the same four conditions when aligned to saccade onset (Figures 3C and 3D; p < 0.001; Table S7). Here, firing increased for good-object saccades but decreased for bad-object saccades, indicating a strong value-based modulation of saccade initiation. This pattern aligns with the notion that Cluster 2 integrates object valence into oculomotor execution.

In contrast, Cluster 3 generally decreased its activity after saccade onset (Figures 3E and 3F). A parametric bootstrap test confirmed significant differences among the four conditions (p < 0.001; Table S8), with especially pronounced suppression observed during bad-object saccades. Hence, Cluster 3 may exert a modulatory influence that is more closely associated with suppressing unwanted movements or signaling aversion-related processing.

### Correlation between GPe neuronal activity and saccade RT

Herein we tested whether the timing of neuronal modulation correlates with saccade initiation, providing insight into whether GPe activity directly influences the timing of movement execution.

To assess whether changes in GPe firing directly influence the speed of saccade initiation, we examined correlations between neuronal activity and saccadic RTs. We focused on trials with contralateral good objects and divided them into four RT groups (Figure 3G).

Clusters 1 and 2 showed early increases in firing on trials with shorter RTs, resulting in median correlation coefficients significantly below zero (Figure 3H; Cluster 1, median = -0.56, p = 3.16 × 10^−6^; Cluster 2, median = -0.24, p = 2.16 × 10^−2^; Wilcoxon signed-rank test). This negative correlation suggests that heightened activity in these clusters promotes faster saccade initiation.

In contrast, the median correlation coefficient for Cluster 3 did not differ significantly from zero (median = 0.12, p = 0.345), implying that this Cluster’s firing rate is not closely tied to saccade onset timing. Hence, Cluster 3 appears less involved in the immediate execution of saccades, reinforcing its potential role in other aspects of the task.

### Distinct GPe neuronal activity patterns during proactive and reactive inhibition

In this section, we contrasted proactive (choice-task) and reactive (fixation-task) inhibition to clarify how Clusters 2 and 3 contribute to suppressing unwanted movements.

Next, we investigated whether the decreased activity of Clusters 2 and 3 in bad-object trials reflected a proactive (goal-directed) form of inhibition, distinct from the reflexive inhibitory demands of a fixation task. Proactive inhibition involves volitional suppression of unwanted actions (26–29), whereas reactive inhibition arises more automatically in response to external cues.

In the choice task, monkeys could reject a bad object with either a “return” saccade or by “staying” in place (Figure 4A). Clusters 2 and 3 showed similarly reduced firing for both rejection strategies (return or stay) in contralateral and ipsilateral directions (Figures 4I, 4L, 4O, and 4R). In contrast, in the fixation task (Figure 4B)—which primarily requires reactive inhibition (30)—monkeys maintained fixation while a good or bad object was presented, and the same two clusters exhibited a markedly smaller reduction in firing when bad objects were shown (Figures 4J, 4M, 4P, and 4S). A quantitative comparison of average normalized firing (100-300 ms post-object) found no significant difference between “return” and “stay” reactions in the choice task (Figures 4K, 4N, 4Q, and 4T; p > 0.05); however, it revealed significant differences between the choice-(return/stay) and fixation-task (good/bad) conditions (p < 0.001; Table S9). Hence, the diminished firing of Clusters 2 and 3 for bad objects is more closely tied to proactive inhibition.

**Figure 4.**
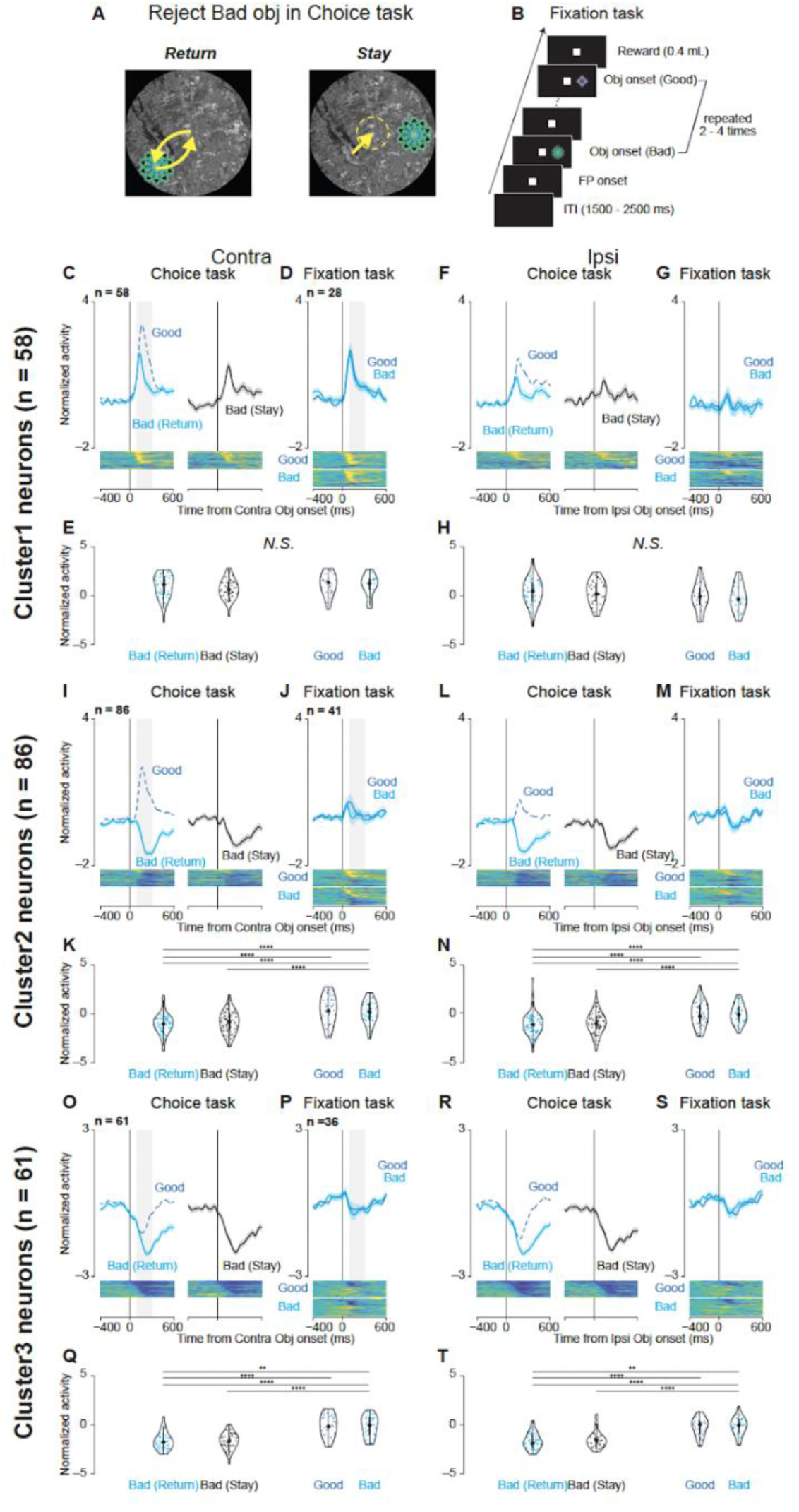
Comparison of neuronal activity during "return" and "stay" responses when rejecting bad objects in the choice and fixation tasks. (A) Schematic illustrations of eye movements during "return" (left) and "stay" (right) responses when rejecting a bad object in the choice task. Yellow arrows indicate saccade directions in the "return" response. The yellow dotted circle indicates that the gaze was maintained around the central fixation point in the "stay" response. (B) Time course of the fixation task. During this task, two objects, one good and one bad, with stable values (same as those used in scene 1 of the choice task) were presented sequentially on either the left or right side of the screen, 2–4 times randomly. The monkeys were required to maintain fixation on the central fixation point throughout the object presentation period to obtain a reward at the end of the trial. (C, F, I, L, O, R) Mean normalized population activity of Cluster 1 (C, F), 2 (I, L), and 3 (O, R) neurons when the monkeys rejected bad objects in scene 1 using the "return" (cyan line) or "stay" (black line) strategy. Data are aligned to the onset of the contralateral (C, I, O) or ipsilateral bad object (F, L, R). The blue dashed line represents the mean normalized population activity when the monkeys accepted good objects in scene 1 for comparison. The shaded areas indicate ± SEM. The lower panels show color maps of the normalized neuronal activity of individual neurons. (D, G, J, M, P, S) Mean normalized population activity of Cluster 1 (D, G), 2 (J, M), and 3 (P, S) neurons during the fixation task when good (blue line) or bad (cyan line) objects were presented on the contralateral (D, J, P) or ipsilateral (G, M, S) side. (E, H, K, N, Q, T) Violin plots showing the distribution of the mean normalized neuronal activity of individual neurons during the choice (E, K, Q) and fixation tasks (H, N, T) for Cluster 1 (E, H), 2 (K, N), and 3 (Q, T) neurons. Neuronal activity was measured for a 200-ms interval beginning 100 ms after object onset (gray rectangle in C, D, I, J, O, and P). The format of the violin plots is the same as that in Figures 2B, 2F, and 2J. The larger circle indicates the median value, the thick vertical line shows the interquartile range (IQR), and the thin vertical line indicates the range from the lower to the upper adjacent values (1.5 x IQR below the first quartile and 1.5 x IQR above the third quartile, respectively). The asterisks indicate significant differences in neuronal activity between conditions (post-hoc pairwise t-tests with Bonferroni correction, *p < 0.05, **p < 0.01, ***p < 0.001, ****p < 0.0001). N.S. indicates no significant difference among conditions (p > 0.05). Abbreviations: SEM, standard errors of the mean.

In contrast, Cluster 1 neurons exhibited increased activity during contralateral object presentations under both the choice and fixation tasks (Figures 4C, 4D, 4F, and 4G). This pattern suggests that Cluster 1’s response may reflect general visual processing or a non-specific inhibitory mechanism, rather than proactive inhibition. Overall, these findings support the notion that Clusters 2 and 3 are specifically related to goal-directed suppression of unwanted actions, while Cluster 1 may reflect a more general visual or reactive process.

### Effects of blocking glutamatergic inputs to the GPe on neuronal activity and behavior

Finally, we use pharmacological manipulations to test whether excitatory inputs drive saccade facilitation in Clusters 1 and 2, thereby confirming a causal role for GPe excitation.

To directly assess whether excitatory inputs underlie the saccade facilitation observed in Clusters 1 and 2, we locally injected a mixture of glutamate receptor antagonists—N-methyl-d-aspartate receptor antagonist (carboxypiperazin-4-propyl-1-phosphonic acid, CPP) and aminomethylphosphonic acid receptor antagonist (2,3-dihydroxy-6-nitro-7-sulfamoyl-benzo (F)quinoxaline, NBQX)—into the caudal-dorsal GPe (cdGPe). This region was chosen because it contained the majority of task-responsive neurons in Clusters 1 and 2 (Figure 2). Three monkeys (Cr, Sp, and Ch) underwent these injections: Monkeys Cr and Sp also participated in neuronal recordings, while Monkey Ch did not but served as an anatomical reference through magnetic resonance imaging (MRI) (Figure S3). Each monkey received five sessions of CPP plus NBQX and five sessions of saline (15 sessions each in total).

Inhibiting these excitatory inputs via CPP plus NBQX significantly prolonged saccade RTs for both contralateral good and bad objects (Figure 5A; p < 0.0001; Table S10), whereas saline had no effect (p > 0.05). For instance, the mean RT for contralateral good objects increased from 193.5 ± 10.3 ms pre-injection to 235.1 ± 15.3 ms post-injection, and for contralateral bad objects from 281.1 ± 34.0 ms to 301.9 ± 51.0 ms.

**Figure 5.**
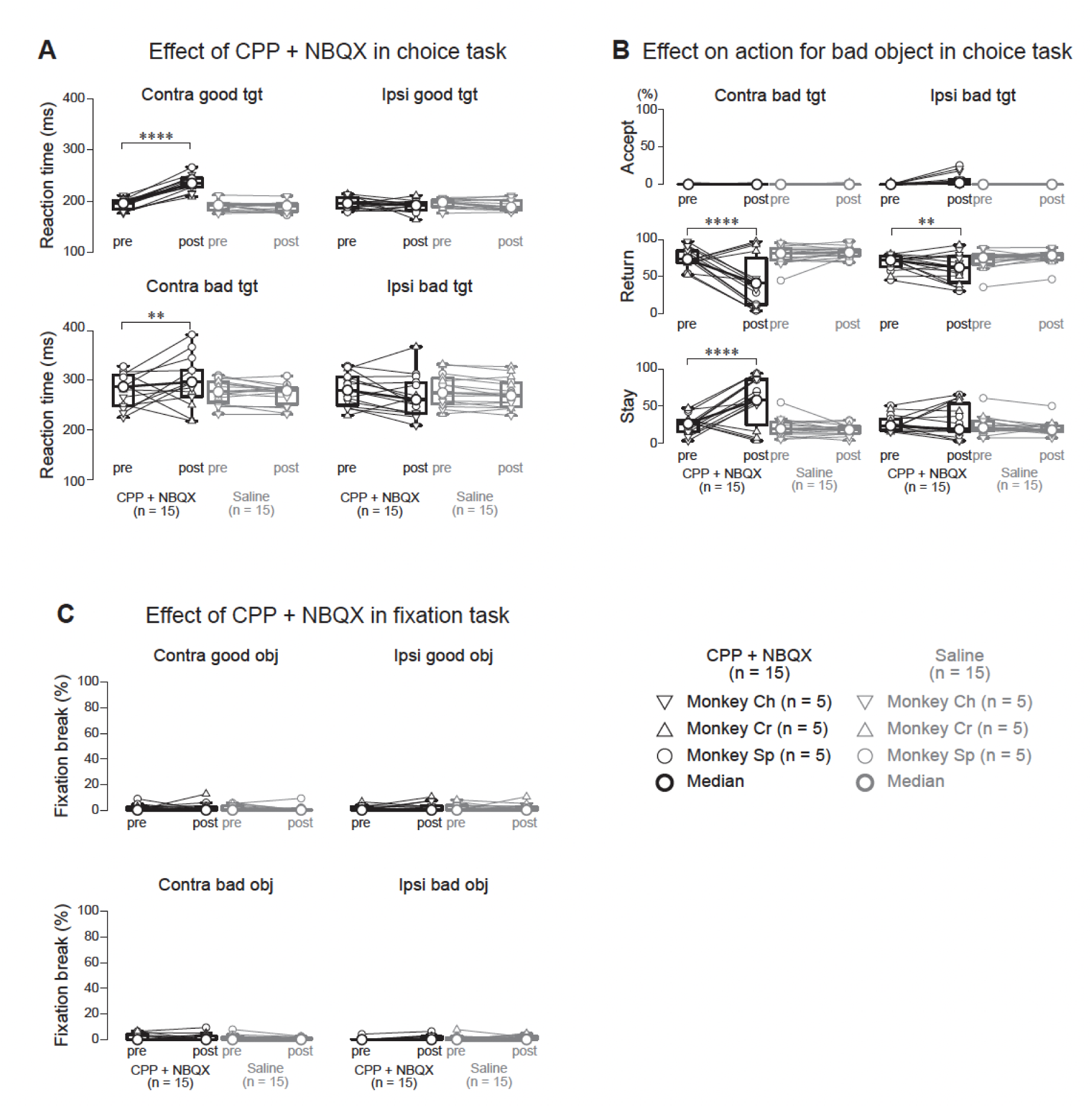
Effects of local injection of glutamatergic receptor antagonists into the cdGPe on behavioral performance during the choice and fixation tasks. (A) Saccadic RTs before (pre) and after (post) local injection of a mixture of CPP and NBQX (CPP+NBQX) or saline into the cdGPe. Data points connected by lines represent median values from individual sessions for each monkey (inverted triangles: Monkey Ch, upright triangles: Monkey Cr, circles: Monkey Sp; n = 5 sessions per condition per monkey). Thick horizontal lines and larger circles indicate population medians. Following CPP+NBQX injection, RTs significantly increased for both contralateral good and contralateral bad objects (p < 0.0001 for both comparisons, parametric bootstrap tests for generalized linear mixed-effects models with Bonferroni correction, see Table S10). In contrast, saline injection had no significant effect on RTs (p > 0.05 for all comparisons). (B) Proportions of different actions (accept, return, stay) for bad objects before (pre) and after (post) CPP+NBQX or saline injection. Data are presented separately for contralateral (left panels) and ipsilateral (right panels) bad objects. Format is the same as in (A). CPP+NBQX injection significantly decreased the proportion of "return" responses and increased the proportion of "stay" responses for contralateral bad objects (p < 0.0001 for both comparisons, parametric bootstrap tests for generalized linear mixed-effects models with Bonferroni correction). A significant, but smaller, decrease in "return" responses was observed for ipsilateral bad objects (p < 0.01). Saline injection had no significant effect on the proportions of "accept", "return", or "stay" responses (p > 0.05 for all comparisons). (C) Proportion of fixation break errors during the fixation task before (pre) and after (post) CPP+NBQX or saline injection. Data are presented separately for contralateral and ipsilateral good and bad objects. Format is the same as in (A). CPP+NBQX injection had no significant effect on the proportion of fixation break errors (p > 0.05 for all comparisons, parametric bootstrap tests for generalized linear mixed-effects models). Asterisks denote significant differences (*p < 0.05, **p < 0.01, ***p < 0.001, ****p < 0.0001, post-hoc pairwise t-tests with Bonferroni correction). Abbreviations: CPP, (±)-3-(2-carboxypiperazin-4-yl) propyl-1-phosphonic acid (N-methyl-d-aspartate receptor antagonist); NBQX, 2,3-dihydroxy-6-nitro-7-sulfamoyl-benzo[f]quinoxaline (aminomethylphosphonic acid receptor antagonist); cdGPe, caudal-dorsal external segment of the globus pallidus; RT, reaction time; tgt, target.

Additionally, CPP plus NBQX injections altered choice behavior for bad objects (Figure 5B). The proportion of “return” responses for contralateral bad objects decreased significantly (p < 0.0001), while “stay” responses increased (p < 0.0001). Saline injections again showed no significant changes (p > 0.05). For contralateral bad objects, the mean “return” proportion dropped from 75.9 ± 14.5% to 43.2 ± 35.9%, whereas “stay” rose from 23.9 ± 14.2% to 56.6 ± 35.7%.

Notably, CPP plus NBQX had no effect on fixation break errors in the fixation task (Figure 5C; p > 0.05), indicating that the observed behavioral alterations were specific to the choice task and did not reflect general motor deficits or changes in reactive inhibition.

Overall, these results demonstrate that the excitatory drive to the GPe— likely from the STN and potentially from cortical or thalamic sources—is essential for normal saccade facilitation in choice tasks. By reducing this excitatory input, CPP plus NBQX likely suppressed the activity of Clusters 1 and 2, thereby prolonging RTs and reducing “return” responses. Therefore, the GPe appears to play a dual role in movement control, supporting suppression of unwanted actions and facilitating goal-directed saccades.

## Discussion

Our findings demonstrate that the GPe plays a dual role in motor control, facilitating desired actions and suppressing unwanted movements. This finding challenges the traditional view of the GPe as merely a homogenous relay in the indirect pathway for motor suppression.

In a novel sequential choice task, we identified three distinct clusters of GPe neurons in macaque monkeys, each exhibiting unique activity patterns associated with saccade facilitation or proactive inhibition. Specifically, Cluster 1 and 2 neurons showed increased activity during the acceptance of good objects with contralateral saccades, reflecting behavioral facilitation. In contrast, Cluster 2 and 3 neurons showed decreased activity when rejecting bad objects, indicating a role in proactive inhibition.

These results highlight the GPe’s functional diversity and underscore a more complex role in motor control than previously recognized. In the following discussion, we first address how GPe activity supports saccade facilitation, followed by its involvement in proactive inhibition.

### Facilitation of saccade generation by increased activity of Cluster 1 and 2 neurons

Our results clearly demonstrate that the increased activity of Cluster 1 and 2 neurons is involved in facilitating saccade generation. This is supported by two main findings. First, the activity of these neurons increased just before saccade initiation towards contralateral good objects (Figures 3A, 3C, and 3E), and the magnitude of this increase was negatively correlated with saccade RTs (Figures 3G and 3H). In other words, the faster the activity increases, the quicker the saccade. Second, local injection of CPP plus NBQX, a mixture of glutamate receptor antagonists, into the cdGPe significantly prolonged saccade RTs for contralateral good and bad objects (Figure 5A). This suggests that the excitatory inputs to the GPe, likely responsible for the increased activity of Cluster 1 and 2 neurons, play a crucial role in facilitating saccade generation.

The origin of these excitatory inputs driving the activity of Cluster 1 and 2 neurons remains to be fully elucidated; however, several lines of evidence point toward the STN as a primary source. Previous studies have demonstrated excitatory projections from the STN to the GPe (31–35), and recent studies in rodents indicate that these projections primarily target prototypical GPe neurons, not arkypallidal neurons (12, 36, 37). Given the electrophysiological characteristics of our identified clusters, particularly their relatively high baseline firing rates (Figure S2), Cluster 1 and 2 neurons are likely predominantly prototypical. This suggests that the STN is a major source of excitatory drive during saccade facilitation.

However, we cannot exclude the possibility of cortical contributions, especially considering the cognitive demands of our task. While direct cortical projections to the GPe have been identified (13–15), these primarily target arkypallidal GPe neurons in rodents, and their functional role in primates, particularly in the context of saccade facilitation, remains unclear. Therefore, although the STN is a likely primary source of excitation, a contribution from cortical inputs cannot be ruled out. For instance, cell-type-specific optogenetic manipulation would allow us to selectively activate or inhibit these inputs to the GPe while recording from identified GPe neuron clusters, thereby testing the causal influence of each pathway on GPe activity during behavior. Additionally, pathway-specific electrophysiology (e.g., antidromic activation) could help differentiate responses evoked by the STN versus cortical inputs based on their latency and synaptic properties.

By systematically employing these approaches we can gain a more precise understanding of the circuitry underlying GPe-mediated saccade facilitation and its implications for motor control and decision-making.

### Involvement of the GPe in proactive inhibition

In addition to its role in saccade facilitation, our results also suggest that the GPe, particularly through the activity of Cluster 2 and 3 neurons, is involved in proactive inhibition. These neurons exhibited decreased activity during the rejection of bad objects, regardless of the specific rejection strategy employed ("return" or "stay"; Figures 4I and 4O). Importantly, this activity decrease was significantly attenuated during the fixation task, which primarily engages reactive inhibition (Figures 4J and 4P). Such a dissociation strongly suggests that the decreased activity of Cluster 2 and 3 neurons is more closely related to proactive inhibition—the volitional, goal-directed suppression of unwanted actions—than to reactive inhibition, which is typically reflexive and triggered by external stimuli (26).

This interpretation aligns with the established role of the indirect pathway in proactive inhibition, where inhibitory projections from the striatum to the GPe suppress GPe activity. Furthermore, our findings are strikingly consistent with those of recent studies demonstrating the involvement of the anterior striatum in proactive inhibition (30). In our previous study, we identified a cluster of neurons in the anterior striatum of macaques performing the same choice and fixation tasks. These neurons increased their activity during the rejection of bad objects. Given the inhibitory nature of striatal projections to the GPe, the increased activity of striatal Cluster 2 neurons would be expected to lead to the decreased activity observed in GPe Cluster 2 and 3 neurons in our study. This reciprocal pattern of activity strongly suggests a functional connection between anterior striatal Cluster 2 and GPe Clusters 2 and 3 in mediating proactive inhibition.

However, while this functional connection is highly suggestive, it is important to note that direct anatomical connectivity between these specific neuronal populations has yet to be established. Future studies employing techniques such as antidromic stimulation, optogenetics (combined with cell-type specific promoters), or retrograde tracing (combined with in situ hybridization) are needed to confirm this connection and elucidate the precise circuitry underlying proactive inhibition within the basal ganglia.

### Possible roles of Cluster 1 neurons: visual responses or reactive inhibition

In contrast to Clusters 2 and 3, Cluster 1 neurons showed increased activity during the presentation of contralateral bad objects in both the choice and fixation tasks (Figures 4C, 4D, 4F, and 4G). While this increased activity could be interpreted as being related to reactive inhibition, the results of the CPP plus NBQX injection experiments suggest otherwise. If the increased activity of Cluster 1 neuron activity during the presentation of contralateral bad objects were primarily related to reactive inhibition, we would expect to see increased fixation break errors during the fixation task following CPP plus NBQX injection, as the blockade of excitatory inputs would impair the ability to suppress reflexive saccades. However, no such increase was observed (Figure 5C). This finding strongly suggests that the increased activity of Cluster 1 neuron activity during bad object presentation is not primarily driven by reactive inhibition.

Considering their strong direction selectivity, increased activity at target onset (Figure 2C) – likely reflecting visual processing –, and the lack of effect of CPP plus NBQX on fixation break errors, we propose that the increased activity of Cluster 1 neuron activity during the presentation of contralateral objects, particularly bad objects, primarily reflects a visual response to the stimuli.

Furthermore, Cluster 1 neurons exhibited even greater activity when contralateral good objects were presented and accepted in the choice task (Figures 2C and 2D). This difference in activity between good and bad object trials suggests that Cluster 1 neurons contribute to saccade generation, with the greater activity for good objects facilitating saccade initiation. The delayed saccade RTs for contralateral good and bad objects following CPP plus NBQX injection in the choice task (Figure 5A) could potentially be explained by a delayed or weakened visual input to the saccade generation system due to the blockade of excitatory inputs to the GPe, which would indirectly affect the downstream targets of Cluster 1 neurons.

Further studies, potentially combining cell-type-specific recording with optogenetic manipulation, could help to definitively disentangle the visual and motor contributions of Cluster 1 neuron activity and clarify the precise role of these neurons in saccade generation and visual processing.

### Comparison with previous studies and implications for the functional organization of the GPe

Our findings extend the conventional view of the GPe as a relay station in the indirect pathway primarily involved in motor suppression (2, 4). While our results are consistent with the role of the GPe in motor inhibition, they also demonstrate that the GPe plays an active role in facilitating movement. This dual role in both movement facilitation and inhibition is in line with recent studies in rodents showing that GPe neurons are heterogeneous and can be classified into distinct populations based on their molecular, anatomical, and physiological properties (6–8, 5). Although we were unable to classify GPe neurons based on cell type-specific markers in this study, the identification of three functionally distinct clusters suggests that a similar heterogeneity may exist in the primate GPe as well.

### Limitations and future directions

This study had a few limitations that should be considered when interpreting the results. First, due to the lacking cell-type-specific markers, we could not definitively link the identified neuronal clusters to specific, genetically defined GPe cell types. This limited our ability to attribute the observed functions (saccade facilitation and proactive inhibition) to particular GPe cell types; hence, future studies employing techniques like optogenetics or DREADDs (Designer Receptors Exclusively Activated by Designer Drugs) are warranted. For example, the possibility remains that different cell types within a single cluster have distinct projection patterns and contribute differentially to behavior.

Second, while our results and previous studies strongly suggest the STN as the primary source of excitatory input driving Cluster 1 and 2 activity, we cannot exclude contributions from other sources, such as the cortex. This uncertainty limits our understanding of the complete circuit mechanisms underlying saccade facilitation. Pathway-specific tracing and manipulation techniques are needed to resolve this.

Third, our pharmacological inactivation targeted the entire cdGPe. Although we focused on this region based on our data recordings, we cannot rule out the involvement of other GPe neurons, or even neighboring regions, in the observed behavioral effects. This could mean that the role of specific clusters is either over- or underestimated. Hence, cell-type-specific manipulations and more localized targeting are necessary for future studies.

Addressing these limitations will require the development and application of precise techniques for manipulating and recording specific cell types and pathways within the basal ganglia. Such approaches are crucial for establishing causal links between neural activity and behavior, ultimately enabling a more complete understanding of basal ganglia function.

### Conclusion

In conclusion, this study provides new evidence that the GPe is crucial for both behavioral facilitation and inhibition. The increased activity of Cluster 1 and 2 neurons appears to be especially important for generating saccades, whereas the decreased activity of Cluster 2 and 3 neurons may underpin proactive inhibition. These findings highlight the GPe’s functional diversity and suggest that it plays a more complex role in motor control than previously recognized. Elucidating the specific circuit mechanisms and cell types within the GPe could further our understanding of basal ganglia disorders and pave the way for targeted therapeutic interventions.

## Methods

The behavioral tasks and experimental procedures in the present study largely followed those established in our previous study on striatal activity (30). We outline the key procedures below, with further details available in our prior publication.

### Animal preparation

All experimental procedures adhered to the Public Health Service Policy on Laboratory Animal Care and were approved by the National Eye Institute Animal Care and Use Committee. Three male rhesus macaques (Macaca mulatta, 8–10 kg), designated as Monkeys Ch, Cr, and Sp, served as subjects. These animals were also utilized in our previous study (30). Surgical implantation of a plastic head holder and recording chambers was performed under isoflurane anesthesia and aseptic conditions. Following recovery, the monkeys underwent training on oculomotor tasks. During experiments, head position was fixed and eye movements were monitored at 1000 Hz using an infrared eye-tracking system (EyeLink 1000, SR Research). Fluid intake was controlled to ensure motivation during the tasks. Comprehensive details on surgical and postoperative care are provided in our previous report (30).

### Behavioral tasks

Experiments were performed in a light- and sound-attenuated room. Visual stimuli were displayed via an LCD projector (PJ658, ViewSonic) onto a screen. A custom software package (Blip; http://www.robili.sblip/) controlled task execution and data acquisition. Two behavioral paradigms, a choice task and a fixation task, were employed, consistent with our previous study (30).

### Choice task

The choice task’s structure and timing were consistent with our prior work. To mitigate potential biases, one of six scene-object sets was randomly selected for each session (Figure S1). Each set comprised four scenes, each containing two fractal objects: a "good" object associated with reward, and a "bad" object yielding no reward. Object-reward contingencies remained constant in scenes 1 and 2 but were reversed in scenes 3 and 4. This value-switching design in scenes 3 and 4 served to decouple neural responses related to object value from those driven by visual features and allowed us to examine how GPe neurons represent and update value in a context-dependent manner.

Each trial began with scene presentation (1000 ms) followed by a central fixation period (700 ms). Subsequently, either the good or bad object was presented at one of six peripheral locations (15° eccentricity). The monkey could accept the object by executing a saccade to it and maintaining fixation for at least 400 ms. Rejection was indicated by one of three actions: a brief saccade to the object followed by a return to central fixation ("return"), sustained central fixation ("stay"), or an eye movement away from both the object and the central fixation point ("other"). Acceptance of the good object resulted in a liquid reward (0.4 mL juice). Following rejection, a new object was presented until acceptance occurred.

### Fixation task

Following the identification of task-modulated neurons during the choice task, a fixation task was administered to investigate GPe activity during saccade suppression. During this task, the good and bad objects from scene 1 of the choice task were presented in a sequential manner (2–4 repetitions, 400 ms duration, 400 ms inter-stimulus interval) while the monkey maintained central fixation. Successful maintenance of fixation throughout the entire sequence was rewarded. Saccades directed towards the presented objects were classified as fixation break errors.

### MRI

After implanting recording chambers, an MRI was performed to map cerebral structures and grid apertures. A gadolinium-enhanced contrast medium (Magnevist, Bayer Healthcare Pharmaceuticals) was introduced into the chambers. Recording sites were localized using a 3 Tesla (3T) MRI system (MAGNETOM Prisma; Siemens Healthcare) and three-dimensional (3D) T1-weighted (T1w, MPRAGE) and T2-weighted (T2w, SPACE) sequences, both with a 0.5 mm voxel size.

To improve visualization of the GPe, particularly in younger animals where contrast with surrounding structures can be limited on conventional T1w and T2w images, quantitative susceptibility mapping (QSM) was employed (38–40). QSM images were reconstructed from phase images acquired with a 3D multi-echo gradient echo sequence (repetition time: 50 ms; echo times: 3.7, 10.1, 16.7, 23.4, 30.0, 36.6, 43.2 ms) (40). The QSM reconstruction pipeline included multiple steps: Phase Unwrapping, where phase images were unwrapped; Background Field Removal, where high-pass filtering was applied to the phase images; and Dipole Inversion, where dipole field inversion was performed to generate quantitative susceptibility maps.

These steps were implemented using the morphology-enabled dipole inversion toolbox (http://pre.weill.cornell.edu/mri/pages/qsm.html) (41) in MATLAB 2019 (The MathWorks, Inc., Natick, MA, USA).

To clearly visualize the grid holes in the reconstructed images, we acquired high-resolution T1w anatomical images using a 3D MPRAGE sequence on a Siemens MAGNETOM Prisma 3T scanner. The imaging parameters were as follows: repetition time (TR) = 1640 ms, echo time (TE) = 2.4 ms, field of view (FOV) = 128 × 128 mm², matrix size = 384 × 384, slice thickness = 0.35 mm, resulting in a voxel size of 0.33 × 0.33 × 0.35 mm³. The acquisition time for this sequence was approximately 11 min.

For this scan, the grid holes were filled with an agar solution mixed with a gadolinium-based contrast agent. We created a fusion image of this high-resolution T1w image and either QSM or a conventional 3D T1w image (Figure S3) to aid in the planning of neural activity recordings and local injections.

### Neuronal recording procedure

Neuronal recordings began after the monkeys demonstrated stable performance, achieving at least 90% accuracy in distinguishing good and bad objects across all four scenes. Single-unit activity within the GPe was recorded using tungsten microelectrodes (1-9 MΩ; Frederick Haer & Co.; Alpha Omega Engineering). Electrodes were advanced through stainless steel guide tubes using a hydraulic micromanipulator (MO-973A, Narishige). Neural signals were amplified, passed through a bandpass filter, which only allows signals within a specific frequency range to pass through (0.3-10 kHz; A-M Systems), and digitized at 40 kHz. Custom voltage-time window discrimination software (Blip) was used for online isolation of individual neurons. This involved setting a threshold to exclude low-amplitude noise and configuring inclusion and exclusion windows (graphical boundaries drawn on the computer screen) to select waveforms (the shapes of the electrical signals) based on their morphology (size and shape). Data collection commenced once stable isolation was achieved and the neuron exhibited task-related activity modulation following target onset in the choice task.

To investigate whether the identified GPe neuron clusters exhibited distinct spike waveform characteristics, we performed spike shape analysis on the extracellularly recorded action (Figures S2C and D). For each isolated neuron, spike waveforms were aligned to the trough (minimum voltage). To normalize the amplitude, each spike waveform was standardized by setting the minimum value to -1 and the subsequent maximum value (peak) to 1. The peak-to-peak duration was then calculated as the time interval between the trough and the peak of the standardized spike waveform.

### Procedure for injecting drugs that block glutamate receptors

To suppress excitatory inputs to the GPe, a mixture of the glutamatergic antagonists CPP (C104, Sigma-Aldrich) and NBQX (N183, Sigma-Aldrich) was injected into the cdGPe, the region where task-related neurons were predominantly located. The characteristically high baseline firing rate of GPe neurons facilitated precise localization of the dorsal boundary of the GPe with the electrode tip. Prior to injection, neuronal activity was recorded using a custom injectrode (42).

Monkeys first performed the choice and fixation tasks to collect pre-injection control data. Then, 1 μL of a 5 mM or 10 mM mixture of CPP and NBQX was injected at 0.2 μL/min using a remotely controlled infusion pump (PHD ULTRA, Harvard Apparatus). Concentrations were based on a previous study (43), with higher levels used to assess the effects of excitatory projections on behavior. After injection, monkeys repeated the choice and fixation tasks to evaluate the effects of CPP and NBQX (5-90 min post-injection).

For statistical analyses, data from 60-90 min post-injection of CPP, NBQX, and saline were used. However, for a subset of data from Monkeys Ch and Sp (using 10 mM CPP and NBQX), analysis was conducted at 20 min post-injection due to pronounced effects that precluded further task performance.

### Data and statistical analysis

All behavioral and neurophysiological data were preprocessed using MATLAB 2022b (MathWorks, Natick, MA, USA). The sample size was determined based on previous studies reporting GPe neuronal recordings (Yoshida and Tanaka, 2016).

### Behavior data analysis

In the choice task, saccade onsets were defined as eye velocities exceeding 40°/s within 400 ms of target onset. RTs for good and bad objects (Figure 1D) included only initial saccades toward objects, excluding return movements. RTs were compared across conditions using Welch’s t-tests. Fisher’s exact test was used to compare "stay" responses between stable (scenes 1 and 2) and flexible (scenes 3 and 4) value conditions (Figure 1E). In the fixation task, saccades toward presented objects were classified as fixation break errors.

### Neuronal data processing

Neuronal data were aligned with event initiation (scene, target, and saccade onset). Peristimulus time histograms (PSTHs) were calculated in 1-ms bins and smoothed with a Gaussian filter (σ = 20 ms). Neuronal activity was Z-transformed by subtracting the baseline firing rate (average firing during the 500ms before event onset) from the smoothed PSTH and dividing it by the PSTH’s standard deviation (SD) (44, 45). The time course of responses was analyzed for each condition using these Z-transformed PSTHs.

Neurons were categorized using k-means clustering, a widely used unsupervised machine learning algorithm suitable for partitioning data into distinct groups based on similarity. This method was chosen because it is efficient, scalable, and does not require prior assumptions about the underlying structure of the data, making it appropriate for exploring the potential heterogeneity of GPe neuronal activity. The clustering was performed based on the neurons’ average standardized firing rate during the presentation of contralateral good or bad objects. Neuronal activity was standardized using a Z-transform. Mean activity was calculated within a 200 ms window (100-300 ms post-object onset) for both contralateral good and bad object presentations.

To determine the optimal number of clusters (K), we employed the silhouette method, a well-established technique for evaluating the quality of clustering. The silhouette value for each neuron measures how similar that neuron is to its own cluster (cohesion) compared with other clusters (separation). The silhouette value ranges from -1 to +1, with higher values indicating better clustering. We simulated silhouette values 5000 times for different numbers of clusters (up to K=6), and the K yielding the highest average silhouette value was selected as the optimal number of clusters. A one-way analysis of variance (ANOVA) followed by post-hoc pairwise t-tests with Bonferroni correction was used to validate the optimal K value (Figure S2A). The Bonferroni-corrected significance level for the post-hoc tests was an α-value of 0.005 (0.05/10).

To assess the relationship between neuronal activity and saccade facilitation, we performed the following analysis (Figures 3G and H). For each neuron, trials were divided into four groups based on saccade RTs within each condition (contralateral/ipsilateral and good/bad object). Groups were ordered by increasing latency (Group 1 = shortest, Group 4 = longest). Mean neuronal activity was calculated for each group within a 200 ms window (100-300 ms post-object onset) and Z-transformed. Correlation coefficients were computed for each neuron to quantify the relationship between group order (saccade latency) and mean neuronal activity. Finally, Wilcoxon signed-rank tests were used to determine if the median correlation coefficient across the population differed significantly from zero.

### Statistical modeling

To compare normalized neuronal activity and behavioral parameters (e.g., saccade reaction times) before and after injections, we used linear mixed-effects models (LMMs) and generalized linear mixed-effects models (GLMMs). These hierarchical models incorporate random effects for individual subjects and neurons, accounting for repeated measurements. By modeling fixed and random effects, LMMs and GLMMs reduce type I errors and better represent the data structure (46). LMMs were used to analyze Z-transformed SNr neuronal activity under various conditions (Figures 2, 3, and S2). GLMMs compared saccade RTs, proportions of chosen actions for bad objects in the choice task, and fixation break errors in the fixation task (Figure 4) pre- and post-injection. For all LMM and GLMM tests, we compared full models with explanatory variables as fixed effects and random effects for monkey, neuron, or session IDs (null model). Detailed model specifications are provided in the Supplementary Methods.

We used a parametric bootstrap method to assess model goodness of fit, performing 10,000 iterations and computing the p-value from the deviance difference between models. If the full model demonstrated significant fit, post-hoc pairwise t-tests with Bonferroni correction were conducted to explore differences. For LMM and GLMM analyses, we used the lme4 (47), pbkrtest (48), emmeans (49), and brms (50) packages in RStudio.

## Resource availability

Further information and requests for resources and reagents should be directed to and will be fulfilled by the lead contact, Atsushi Yoshida.

## Materials availability

This study did not generate new unique reagents.

## Data and code availability

Data and code are available upon request.

## Acknowledgments

This research was supported by the Intramural Research Program at the National Institutes of Health and National Eye Institute (1ZIA EY000415). MRI scanning was conducted in the Neurophysiology Imaging Facility Core (National Institute of Mental Health, National Institute of Neurological Disorders and Stroke, and National Eye Institute). We would like to thank Richard Krauzlis, the lab chief of the laboratory of Sensorimotor Research at the National Eye Institute, for his support as a co-mentor. We would also like to thank D. Parker, H. Warnock, G. Tansey, K. Allen-Worthington, A.M. Nichols, D. Yochelson, J. Fuller-Deets, and M. Robinson for their technical assistance.

## Author contributions

Conceptualization, A.Y.; Software, A.Y.; Formal Analysis, A.Y.; Writing – Original Draft, A.Y.; Writing – Review & Editing, A.Y.; Visualization, A.Y.; Supervision O.H.; Funding Acquisition, O.H.

## Supplementary Methods

### Detailed model specifications

Model specifications for the statistical analyses presented in the main text are detailed below. All linear mixed-effects (LMM) and generalized linear mixed-effects (GLMM) models included random effects (e.g., [1|monkey_ID] or [1|monkey_ID: Neuron_ID]) to statistically control for individual differences at both the subject and neuron levels. By accounting for these hierarchical structures, the models minimized the risk of inflated type I errors due to the nonindependence of observations within the same monkey or neuron, ensuring more accurate and robust inferences.

### 1. Linear mixed-effects model for neuronal activity at scene onset (Figures 2B, G, and K)

Model objective: To evaluate differences in neuronal activity following scene onset.

Model specifications:

Full Model: NormalizedNeuronalActivity ∼ Scene + (1|monkey_ID) + (1|monkey_ID: Neuron_ID)

Null Model: NormalizedNeuronalActivity ∼ (1|monkey_ID) + (1|monkey_ID: Neuron_ID)

Variables:

NormalizedNeuronalActivity: Mean Z-transformed PSTH (100-300 ms after scene onset)

Scene: Fixed effect (levels: 1-4)

monkey_ID, Neuron_ID: Random effects

Statistical threshold: α = 0.05

### 2. Linear mixed-effects model for neuronal activity at target onset (Figures 2D, H, and L)

Model objective: To examine how neuronal activity varies with scene context, object value, and target direction.

Model specifications:

Full Model: NormalizedNeuronalActivity ∼ Scene × Value × Direction + (1|monkey_ID) + (1|monkey_ID: Neuron_ID)

Null Model: NormalizedNeuronalActivity ∼ (1|monkey_ID) + (1|monkey_ID: Neuron_ID)

Variables:

NormalizedNeuronalActivity: Mean Z-transformed PSTH (100-300 ms after target onset)

Scene: Fixed effect (levels: 1-4)

Value: Fixed effect (levels: good, bad)

Direction: Fixed effect (levels: contralateral, ipsilateral)

monkey_ID, Neuron_ID: Random effects

Multiple comparisons:

6 pairwise comparisons between conditions (good vs. bad, contralateral vs. ipsilateral)

Bonferroni-corrected threshold: α = 0.05/6

### 3. Linear mixed-effects model for neuronal activity at saccade onset (Figures 3B, D, and F)

Model objective: To examine neuronal activity patterns aligned to saccade onset.

Model specifications:

Full Model: NormalizedNeuronalActivity ∼ Scene × Value × Direction + (1|monkey_ID) + (1|monkey_ID: Neuron_ID)

Null Model: NormalizedNeuronalActivity ∼ (1|monkey_ID) + (1|monkey_ID: Neuron_ID)

Variables:

NormalizedNeuronalActivity: Mean Z-transformed PSTH (from 150 ms before to 50 ms after saccade onset)

Scene: Fixed effect (levels: 1-4)

Value: Fixed effect (levels: good, bad)

Direction: Fixed effect (levels: contralateral, ipsilateral)

monkey_ID, Neuron_ID: Random effects

Multiple comparisons:

6 pairwise comparisons between conditions (good vs. bad, contralateral vs. ipsilateral)

Bonferroni-corrected threshold: α = 0.05/6

### 4. Linear mixed-effects model for neuronal activity during choice rejection and fixation (Figures 4E, H, K, N, Q, and T)

Model objective: To compare neuronal activity patterns between different rejection strategies and across choice and fixation tasks.

Model specifications:

Full Model: NormalizedNeuronalActivity ∼ Condition × Direction + (1|monkey_ID) + (1|monkey_ID: Neuron_ID)

Null Model: NormalizedNeuronalActivity ∼ (1|monkey_ID) + (1|monkey_ID: Neuron_ID)

Variables:

NormalizedNeuronalActivity: Mean Z-transformed PSTH (100-300 ms post-target)

Condition: Fixed effect (levels: return (choice task), stay (choice task), good (fixation task), bad (fixation task))

Direction: Fixed effect (levels: contralateral, ipsilateral)

Multiple comparisons:

6 pairwise comparisons between conditions (good vs. bad, contralateral vs. ipsilateral)

Bonferroni-corrected threshold: α = 0.05/6

### 5. Generalized linear mixed-effects model for saccade reaction times after injection (Figure 5A)

Model objective: To investigate the effects of glutamatergic antagonist injection on saccadic reaction times.

Model specifications:

Full Model: MedianSaccadeReactionTimes ∼ Injection × PrePost × Value × Direction + (1|monkey_ID) + (1|monkey_ID: Session_ID)

Null Model: MedianSaccadeReactionTimes ∼ (1|monkey_ID) + (1|monkey_ID: Session_ID)

Variables:

MedianSaccadeReactionTimes: Median reaction time per condition Injection: Fixed effect (levels: antagonist, saline)

PrePost: Fixed effect (levels: pre-injection, post-injection)

Value: Fixed effect (levels: good, bad)

Direction: Fixed effect (levels: contralateral, ipsilateral)

Session_ID: Random effect identifying individual injection sessions

Distribution: Poisson

Rationale for distribution choice: Reaction times are non-negative count data characterized by a right-skewed distribution.

Multiple comparisons:

8 pairwise comparisons (pre vs. post for each condition)

Bonferroni-corrected threshold: α = 0.05/8

### 6. Generalized linear mixed-effects model for chosen action rate after injection (Figure 5B)

Model objective: To investigate the impact of glutamatergic antagonist injection on action selection for bad objects.

Model specifications:

Full Model: ChosenActionRate ∼ Injection × PrePost × Value × Direction + (1|monkey_ID) + (1|monkey_ID: Session_ID), weights = (total trial count)

Null Model: ChosenActionRate ∼ (1|monkey_ID) + (1|monkey_ID: Session_ID), weights = (total trial count)

Variables:

ChosenActionRate: Proportion of selected actions

Injection, Pre- Post-, Value, Direction: Fixed effects as described above total_trial_count: Weights to account for different numbers of trials

Distribution: Binomial

Rationale for distribution choice: Analysis of proportional data with binary outcomes

Multiple comparisons:

8 pairwise comparisons (pre vs. post for each condition)

Bonferroni-corrected threshold: α = 0.05/8

### 7. Generalized linear mixed-effects model for fixation break error rate after injection (Figure 5C)

Model objective: To examine how glutamatergic antagonist injection affects the ability to suppress reflexive saccades.

Model specifications:

Full Model: FixBreakErrorRate ∼ Injection × PrePost × Value × Direction + (1|monkey_ID) + (1|monkey_ID: Session_ID), weights = (total trial count)

Null Model: FixBreakErrorRate ∼ (1|monkey_ID) + (1|monkey_ID: Session_ID), weights = (total trial count)

Variables:

FixBreakErrorRate: Proportion of fixation break errors

All other variables, as defined above

Distribution: Binomial

Rationale for distribution choice: Analysis of error rate data with binary outcomes.

Multiple comparisons:

8 pairwise comparisons (pre vs. post for each condition)

Bonferroni-corrected threshold: α = 0.05/8

**Figure S1.**
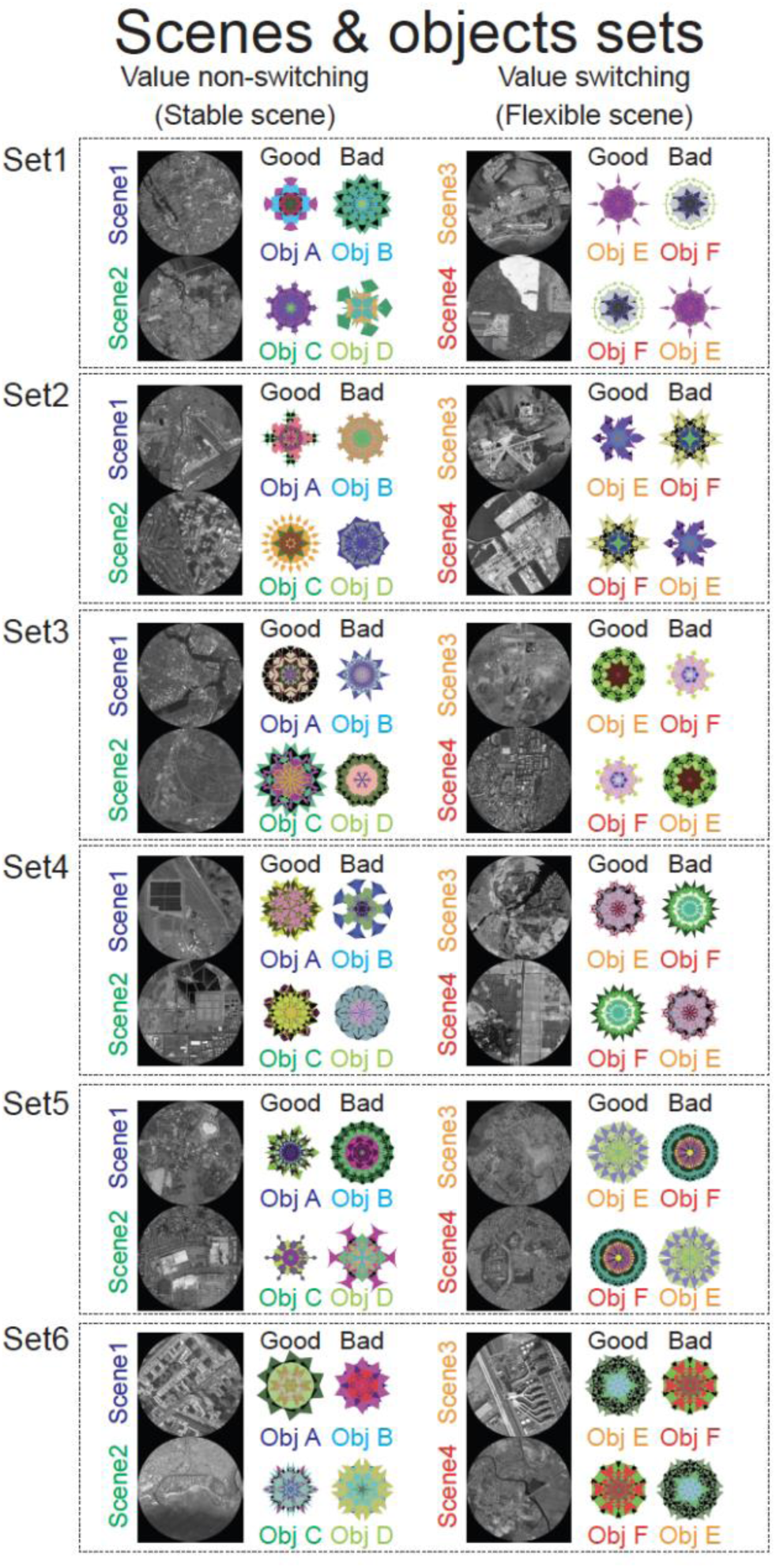
All sets of scenes 1-4 and good and bad objects for the choice task.

**Figure S2.**
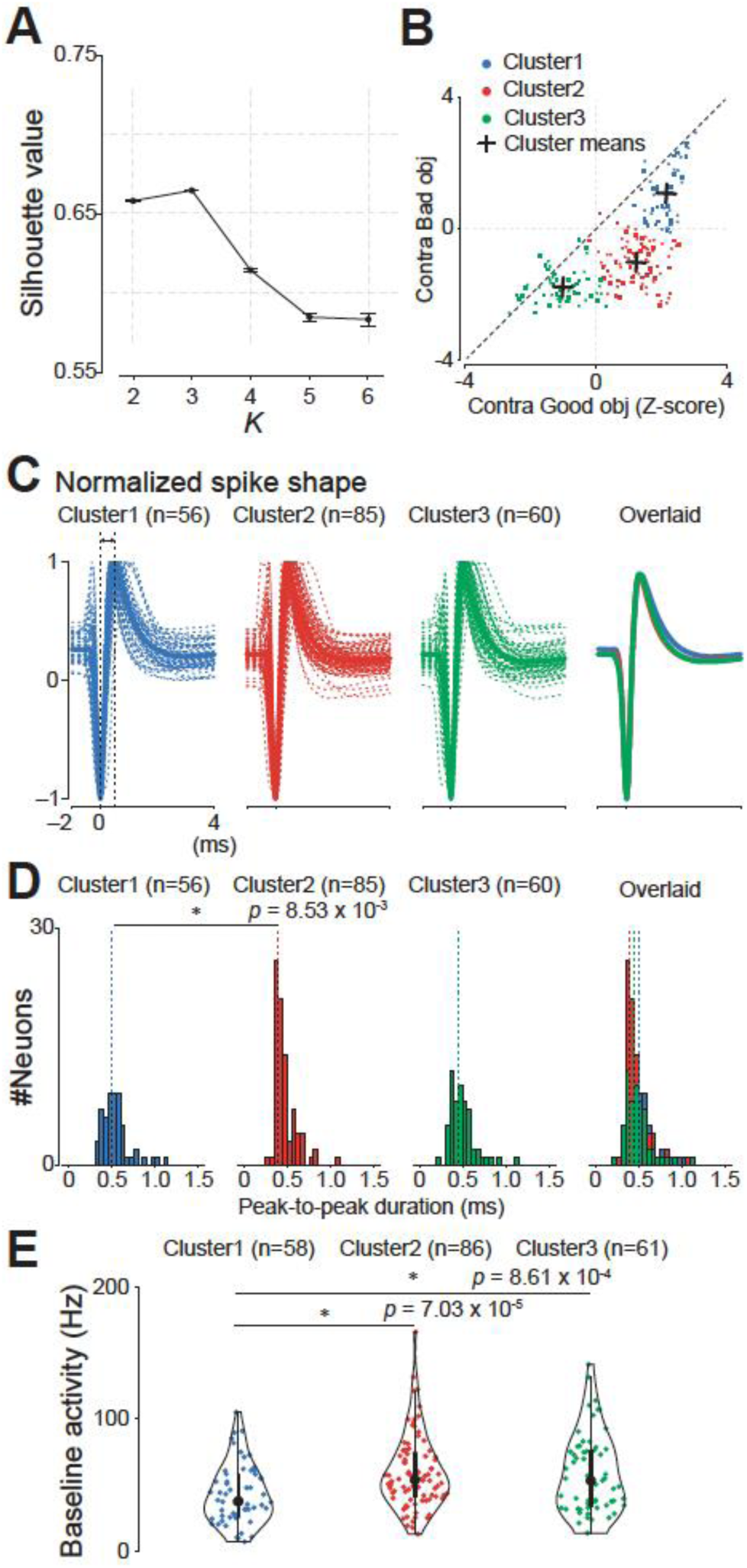
Cluster analysis of GPe neurons based on task-related activity and comparison of their electrophysiological properties. (A) Determination of the optimal number of clusters (K) for k-means clustering. The graph shows the mean ± SD silhouette values for different K values, calculated from 5,000 simulations. The silhouette value was significantly higher for K=3 than for other values (p < 0.001, one-way ANOVA), indicating that the neurons were best classified into three clusters. (B) Scatter plot of the average standardized firing rates of each neuron during the presentation of a contralateral good object (x-axis) and a contralateral bad object (y-axis). The firing rates were calculated for the period between 100 and 300 ms after object presentation and standardized using the z-score method (see Methods for details). Each dot represents a neuron, and the colors indicate the cluster to which the neuron was assigned: Cluster 1 (blue), Cluster 2 (red), and Cluster 3 (green). The crosses indicate the mean values for each cluster. (C) Normalized spike shapes for each cluster. The thick lines represent the average spike shape for each cluster, and the thin lines represent the individual spike shapes. The spike shapes were normalized to the trough and peak amplitudes for each neuron. (D) Histograms of the peak-to-peak duration for each cluster. The peak-to-peak duration was defined as the time between the trough and the peak of the spike waveform. The asterisk indicates a significant difference between Clusters 1 and 2 (p = 8.53 x 10^−3^, Kruskal–Wallis test followed by Dunn’s post-hoc test). (E) Violin plots of the baseline firing rates for each cluster. The baseline firing rate was defined as the average firing rate during the 500 ms period preceding the scene onset. The format of the violin plots is the same as that in Figure 2B. The larger circle indicates the median value, the thick vertical line shows the interquartile range (IQR), and the thin vertical line indicates the range from the lower to the upper adjacent values (1.5xIQR below the first quartile and 1.5xIQR above the third quartile, respectively). The asterisks indicate significant differences between Clusters 1 and 2 (p = 7.03 x 10^−5^) and between Clusters 1 and 3 (p = 8.61 x 10^−4^) (Kruskal–Wallis test followed by Dunn’s post-hoc test). Abbreviations: GPe, external segment of the globus pallidus; SD, standard deviation.

**Figure S3.**
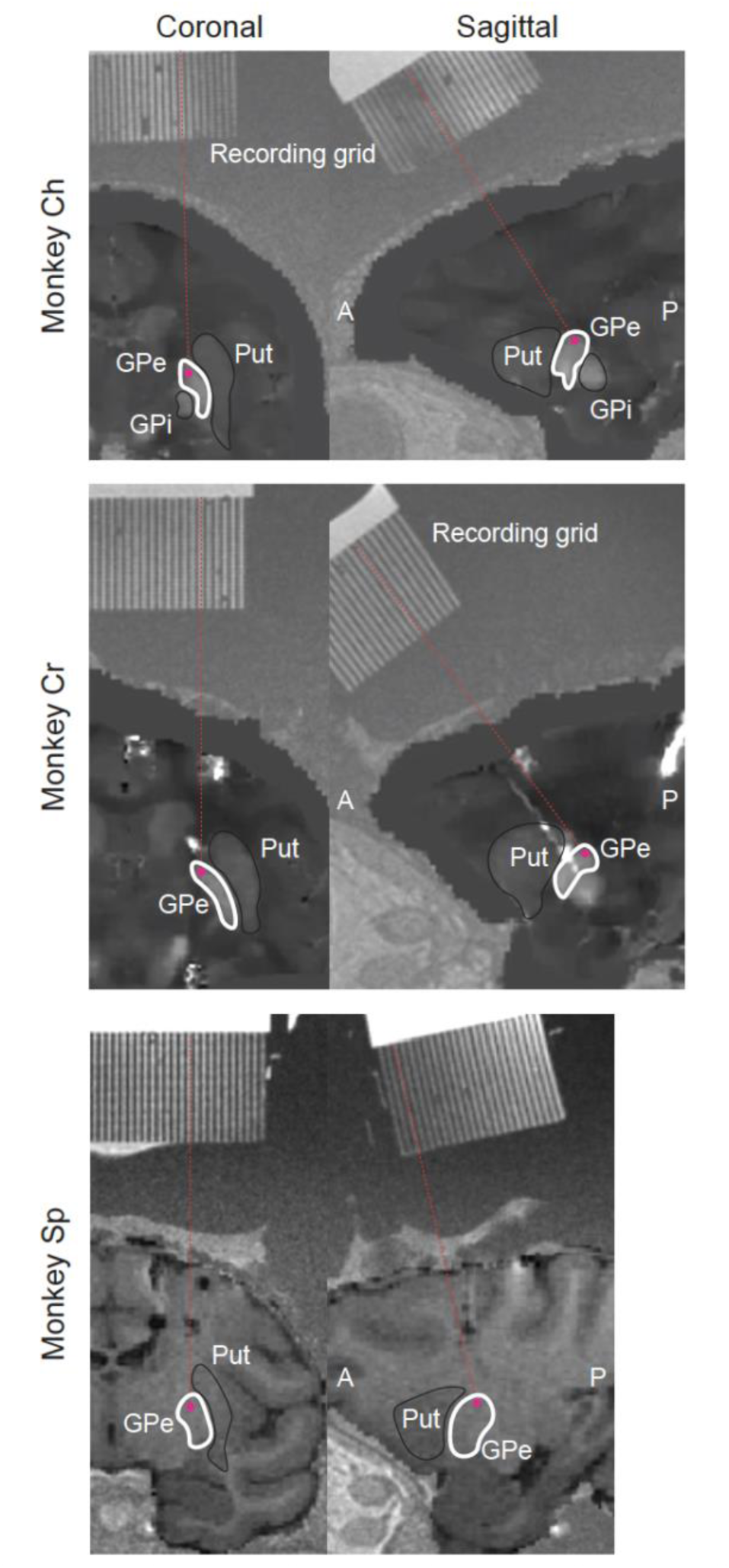
Examples of injection sites in the GPe of three macaque monkeys (Monkey Ch, Monkey Cr, and Monkey Sp). Coronal (left) and sagittal (right) sections of MRI images show representative injection sites (red dots) in each monkey. The MRI images were fused images of high-resolution T1-weighted images and either QSM or a conventional three-dimensional T1-weighted image. The white lines delineate the approximate borders of the GPe and putamen. The red dots indicate the location of the injection site, which was targeted to be in the cdGPe. Abbreviations: A, anterior; P, posterior; cdGPe, caudodorsal globus pallidus external segment; GPe, globus pallidus external segment; GPi, globus pallidus internal segment; Put, putamen; QSM, quantitative susceptibility mapping; MRI, magnetic resonance imaging.

**Table S1.**
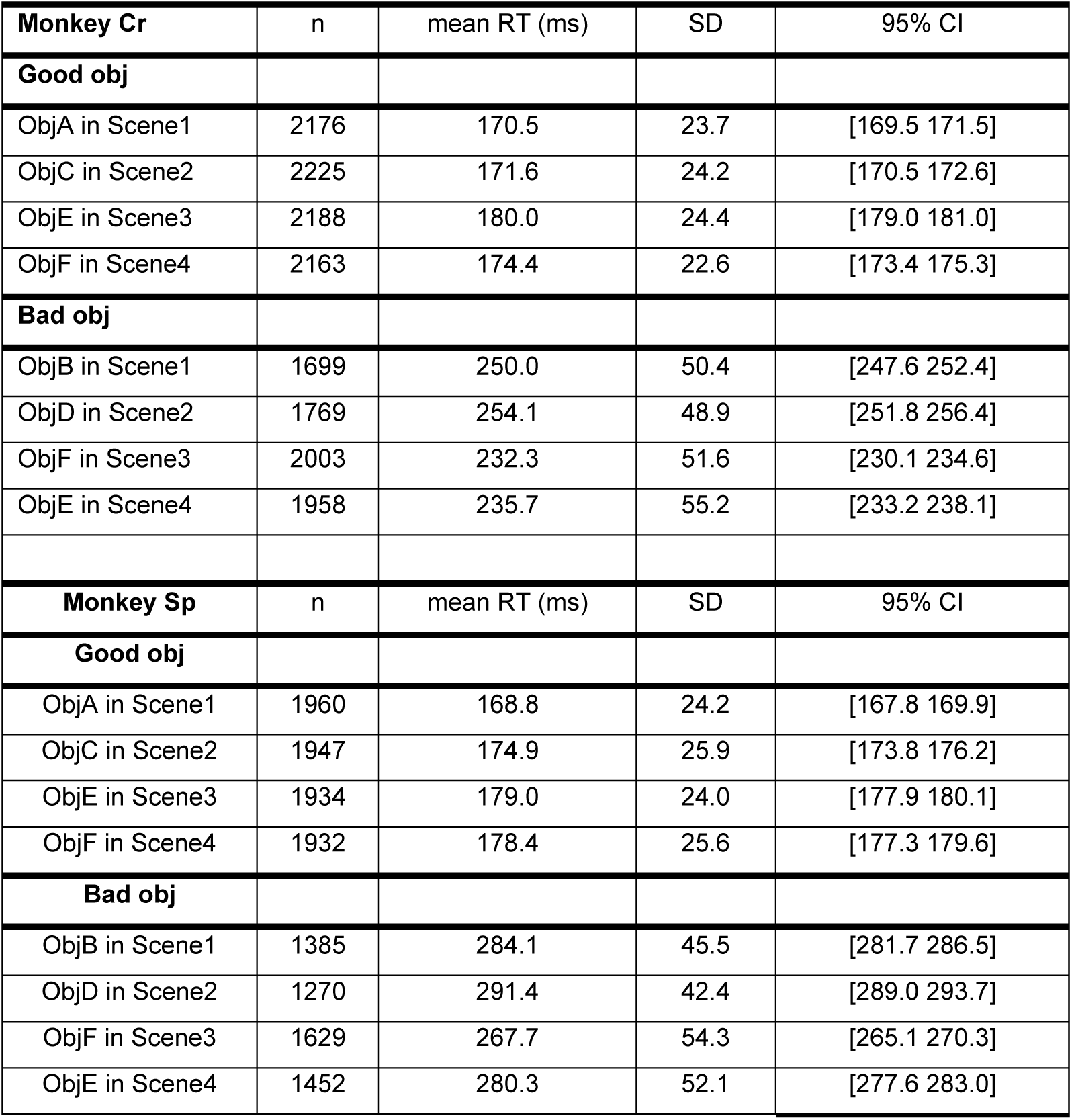
Saccade reaction times in each condition of each monkey.

**Table S2.**
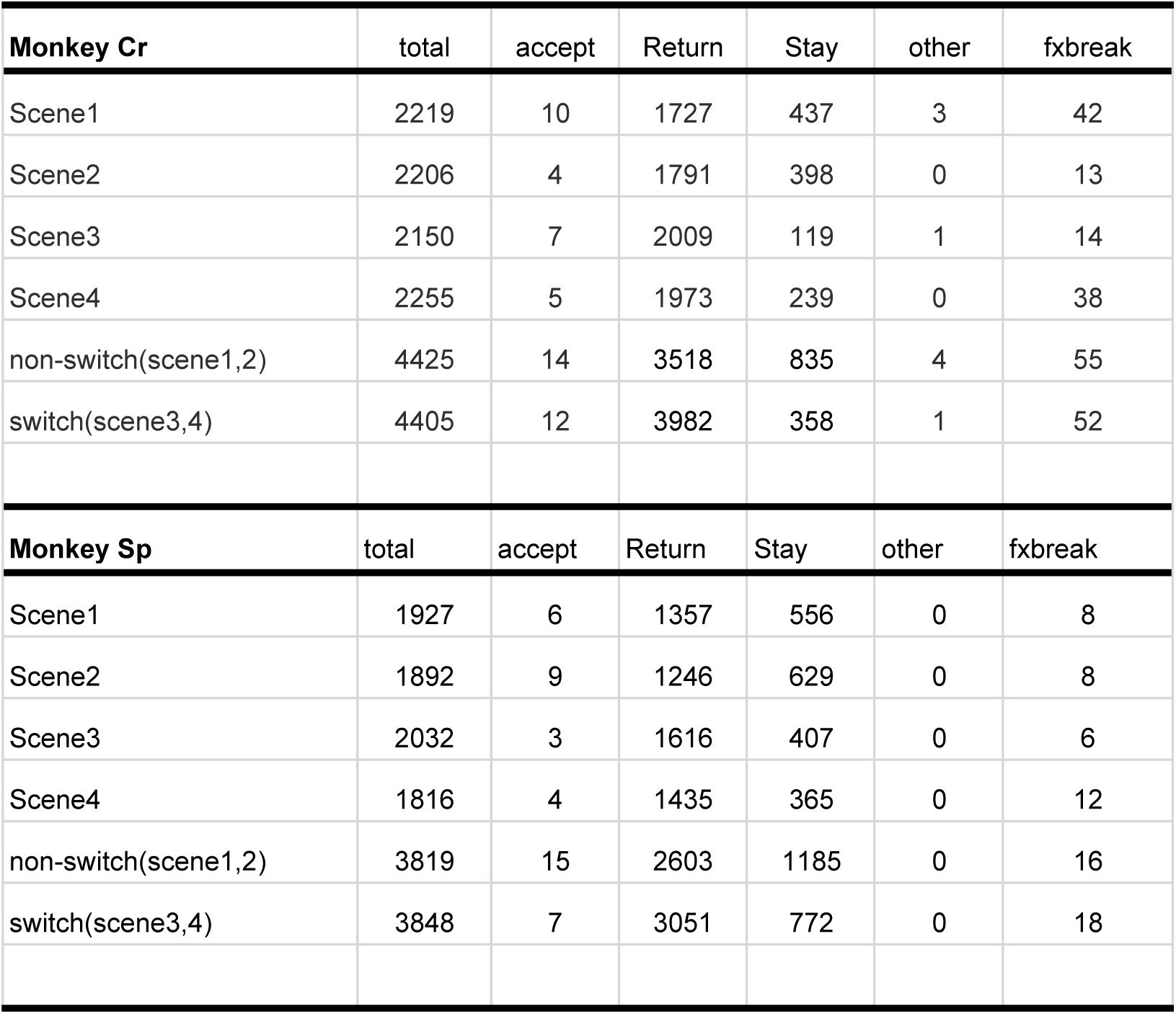
Counts of chosen actions for Bad objects.

**Table S3.**
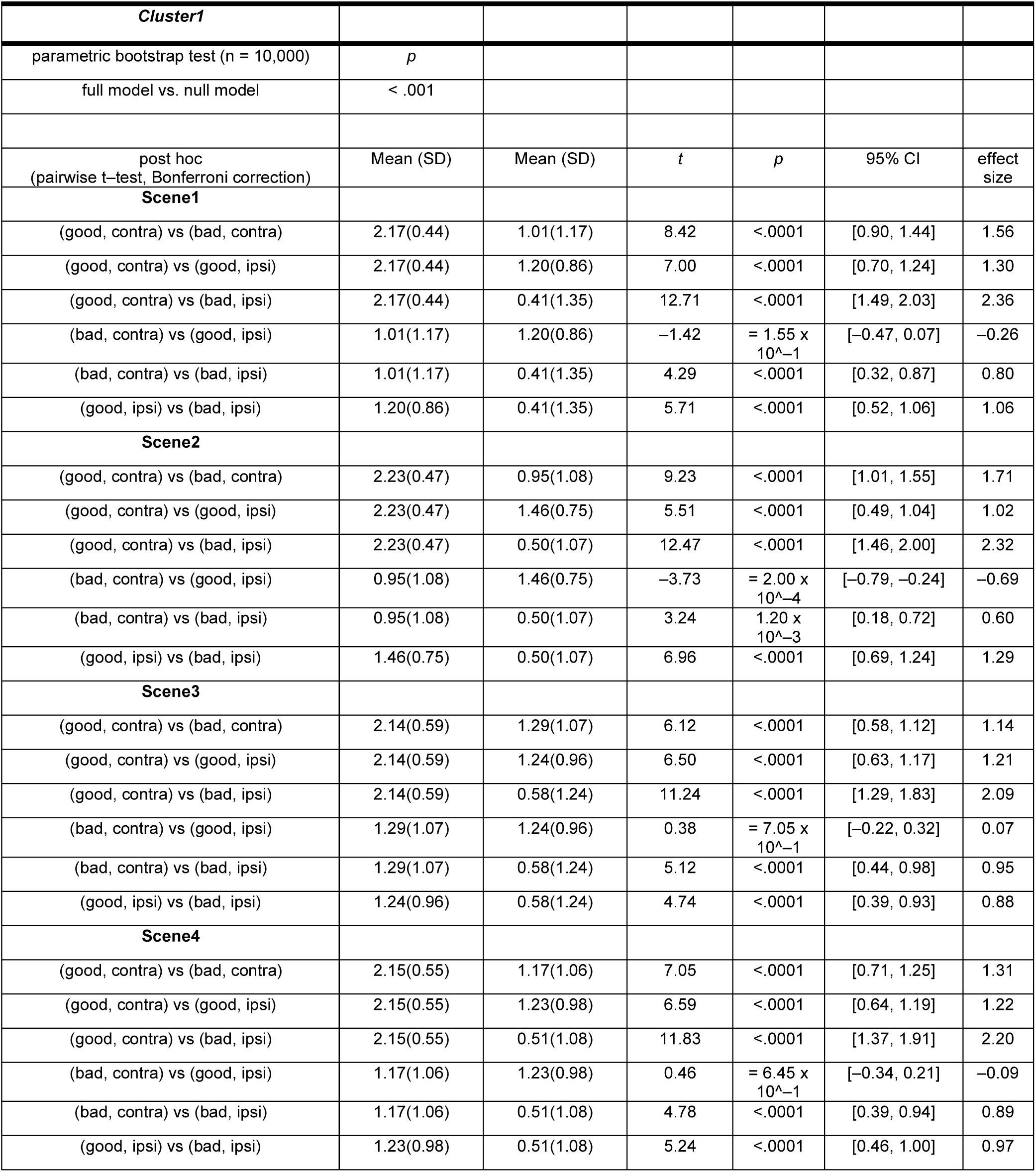
Summary of statistical test to compare the normalized neuronal activity of GPe neurons of cluster1 at target onset among conditions during choice task in Figure 2.

**Table S4.**
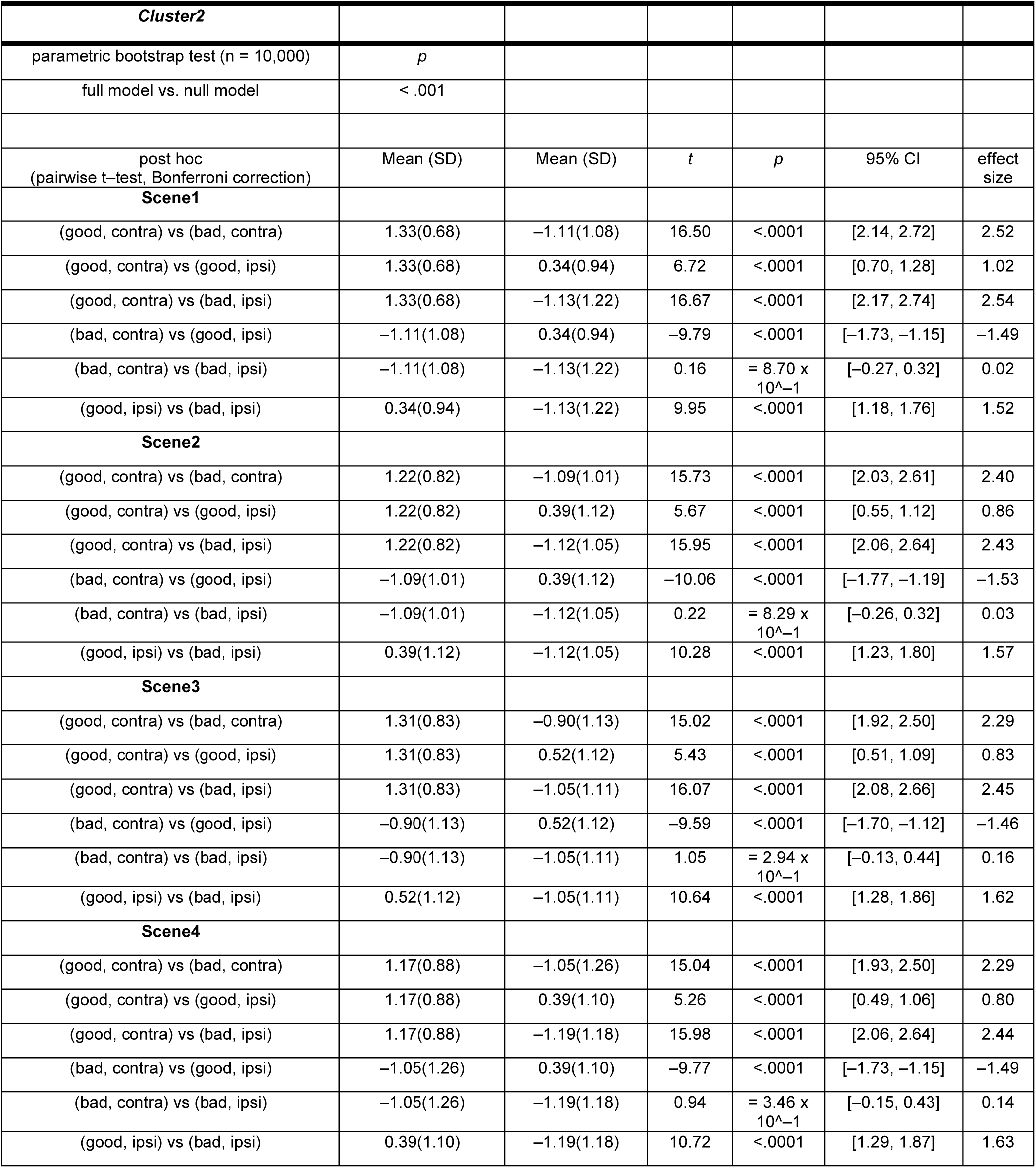
Summary of statistical test to compare the normalized neuronal activity of GPe neurons of cluster2 at target onset among conditions during choice task in Figure 2.

**Table S5.**
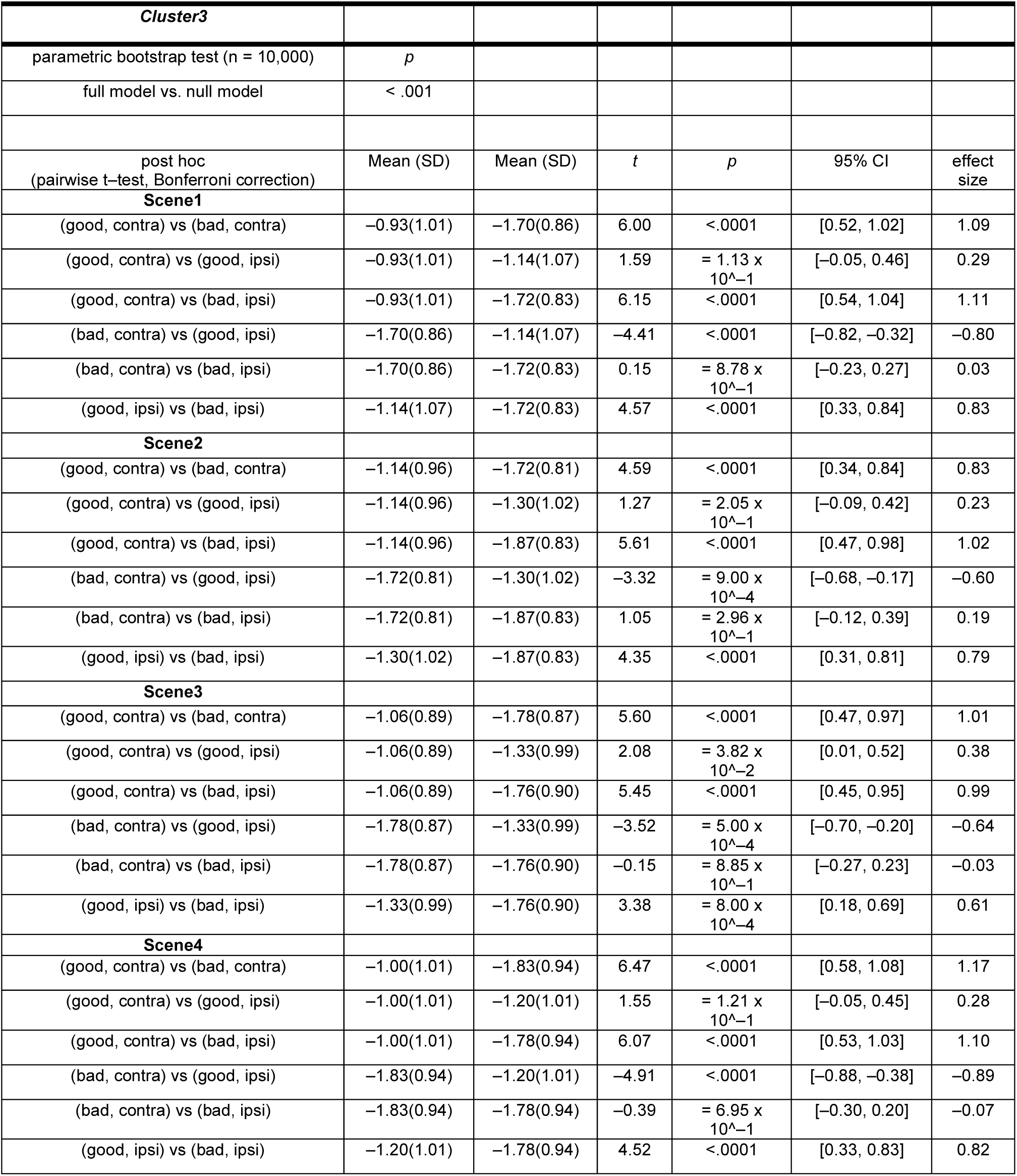
Summary of statistical test to compare the normalized neuronal activity of GPe neurons of cluster3 at target onset among conditions during choice task in Figure 2.

**Table S6.**
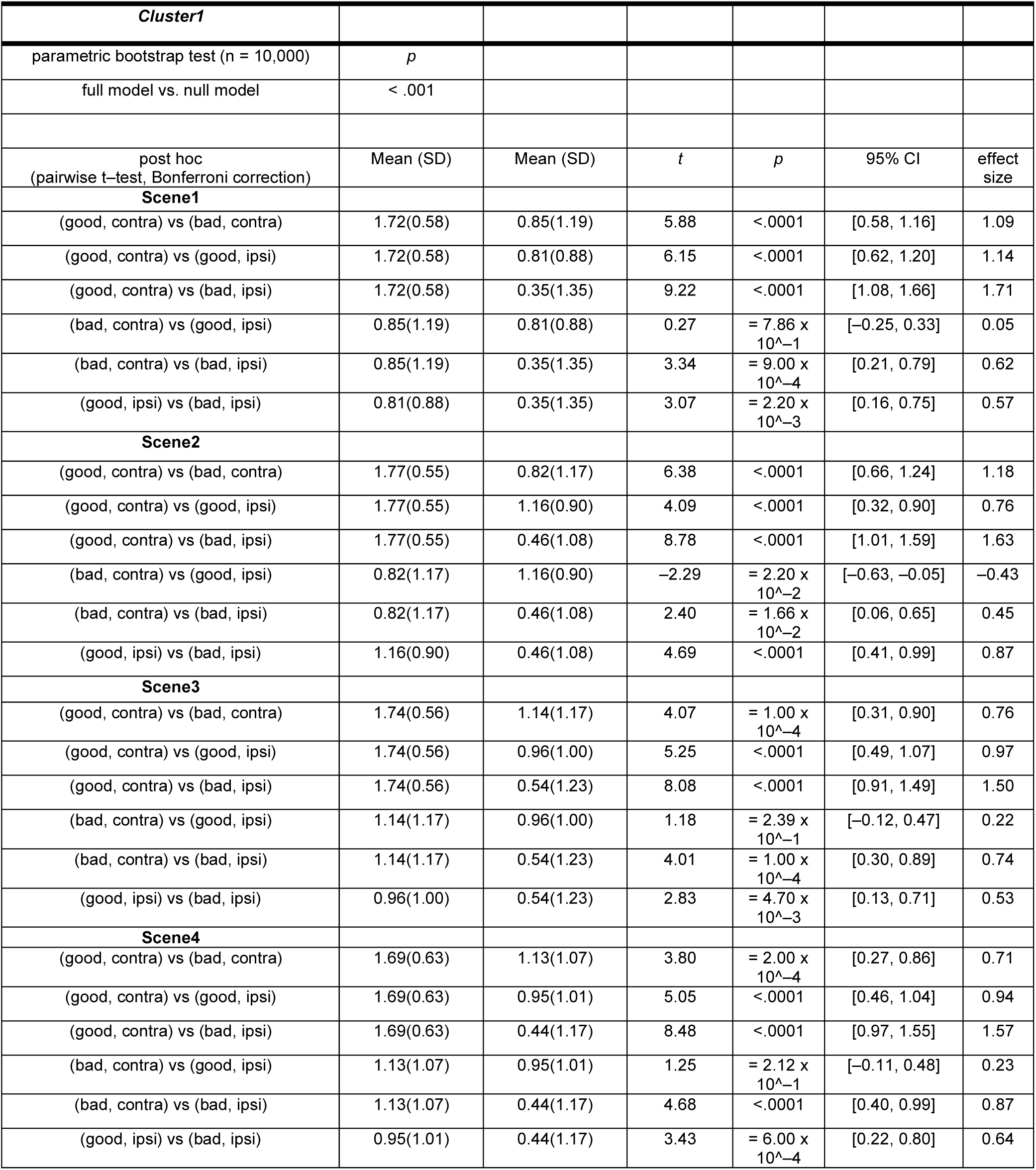
Summary of statistical test to compare the normalized neuronal activity of GPe neurons of cluster1 at saccade onset among conditions during choice task in Figure 3.

**Table S7.**
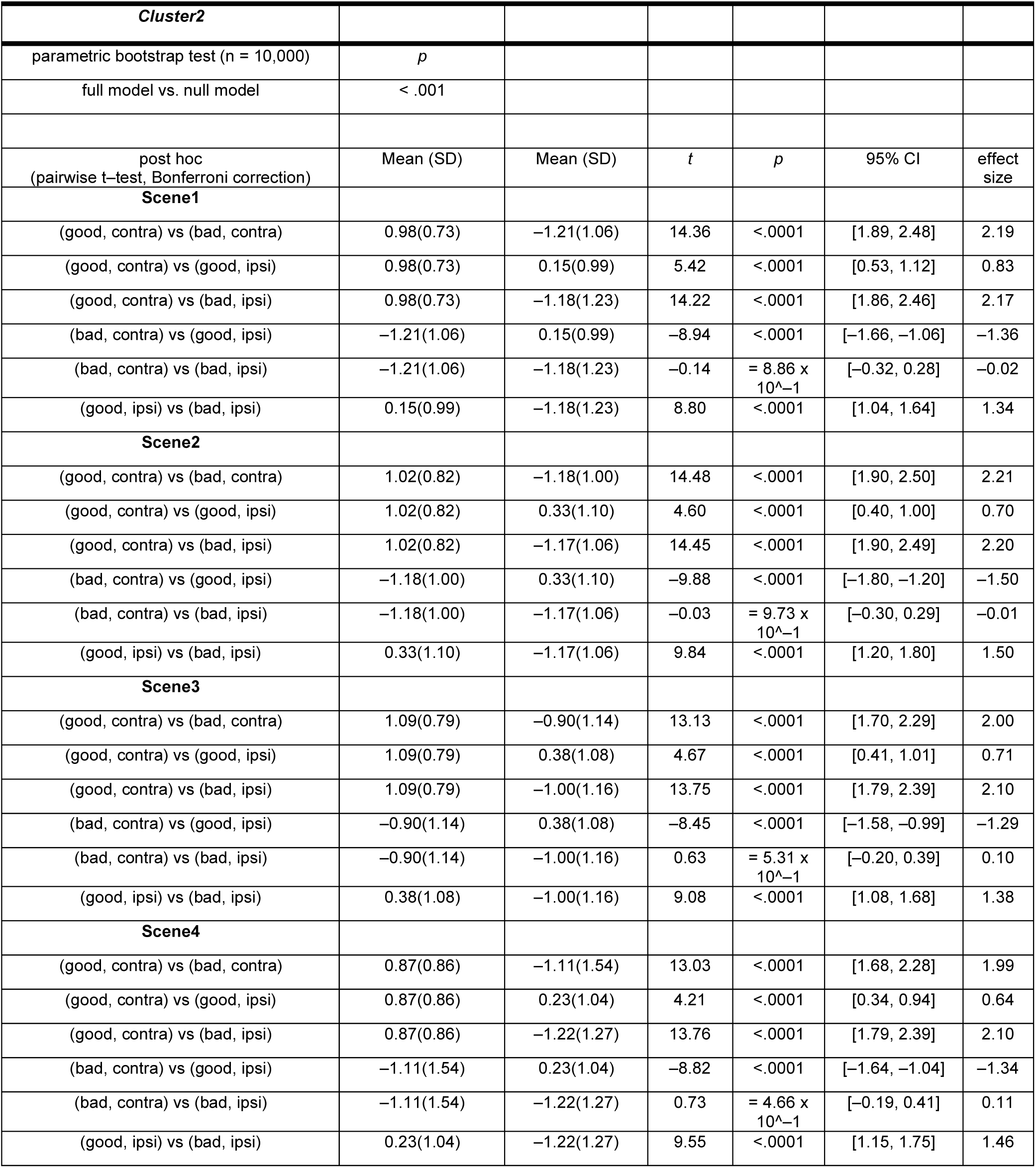
Summary of statistical test to compare the normalized neuronal activity of GPe neurons of cluster2 at saccade onset among conditions during choice task in Figure 3.

**Table S8.**
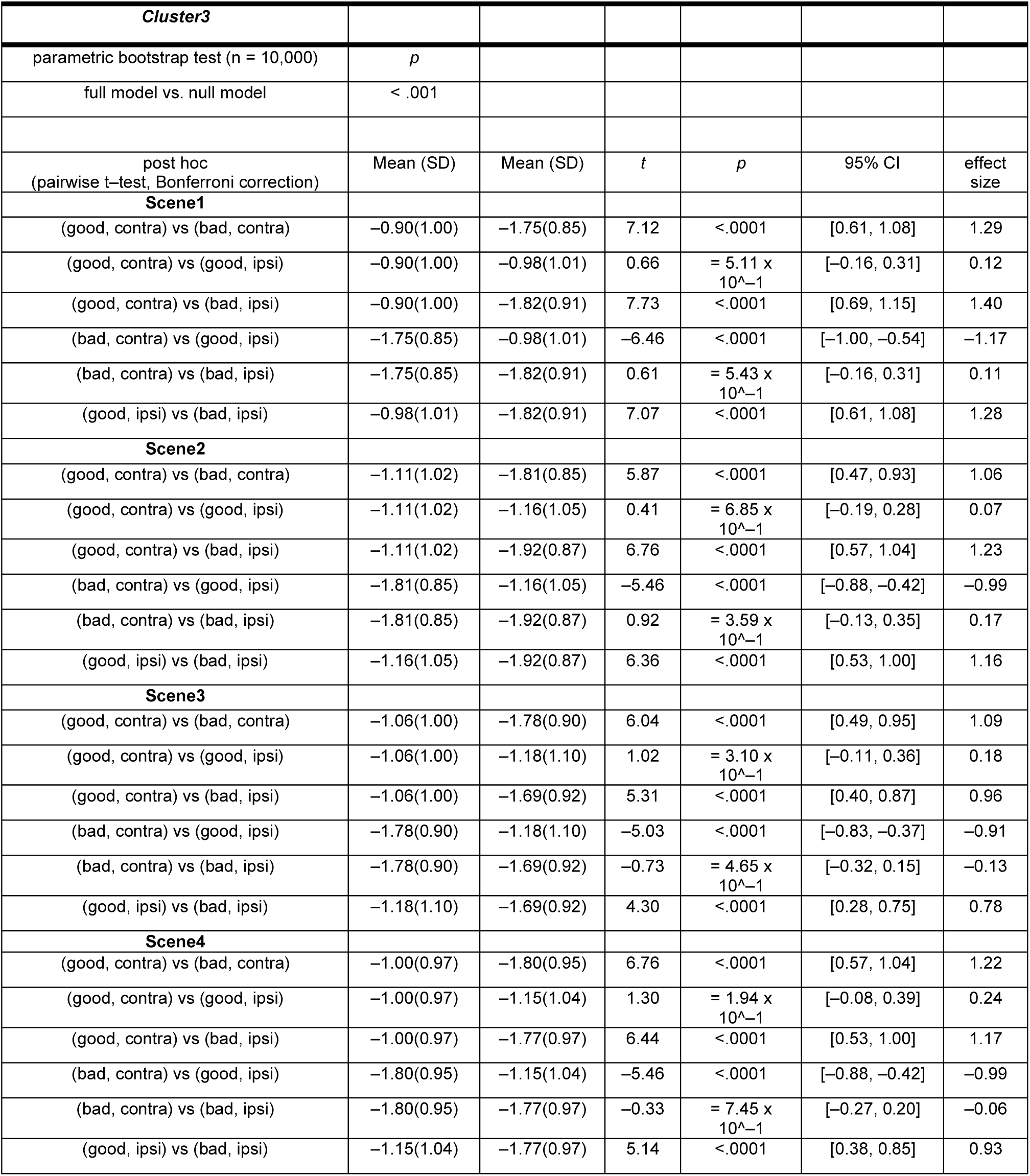
Summary of statistical test to compare the normalized neuronal activity of GPe neurons of cluster3 at saccade onset among conditions during choice task in Figure 3.

**Table S9.**
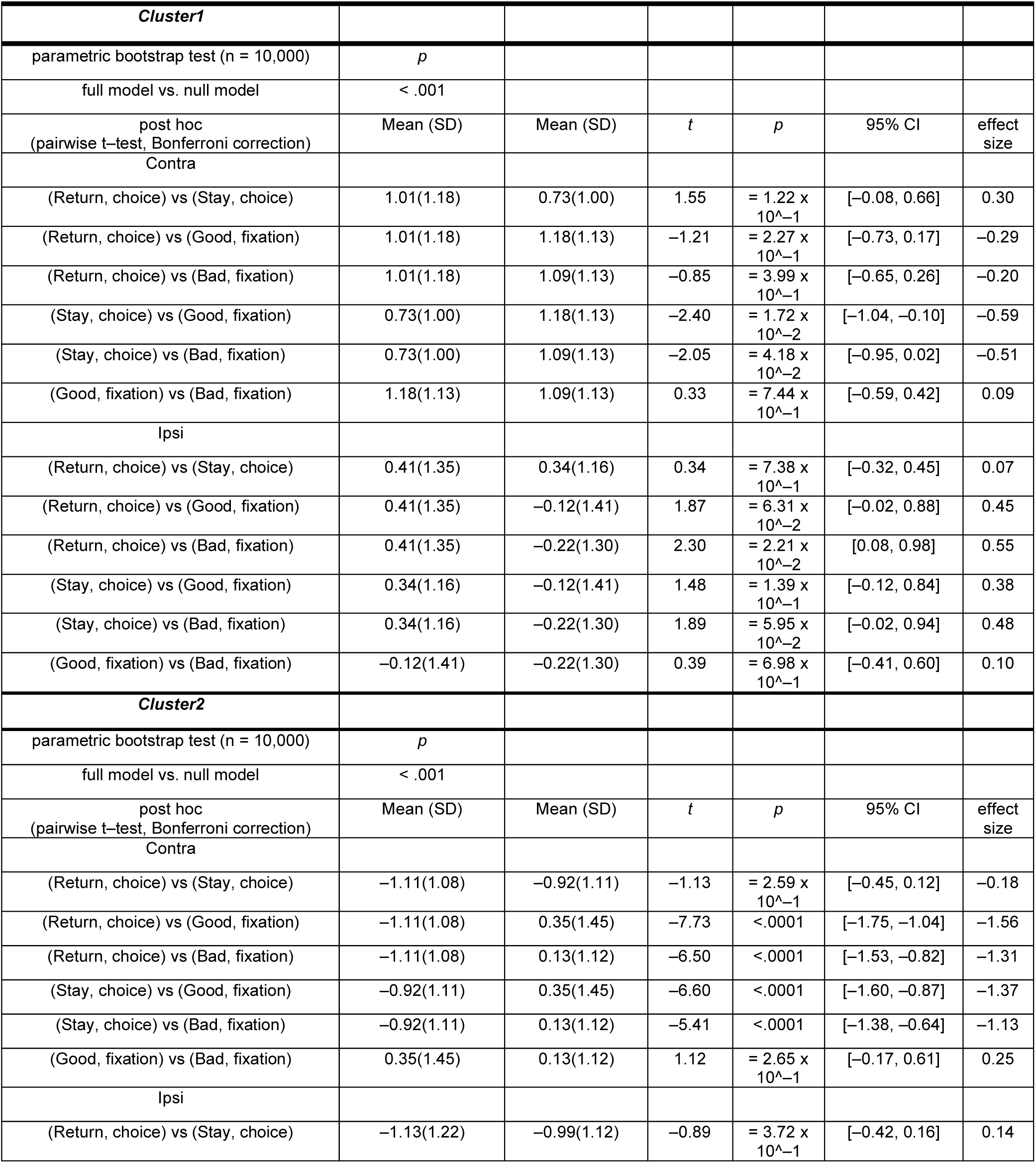
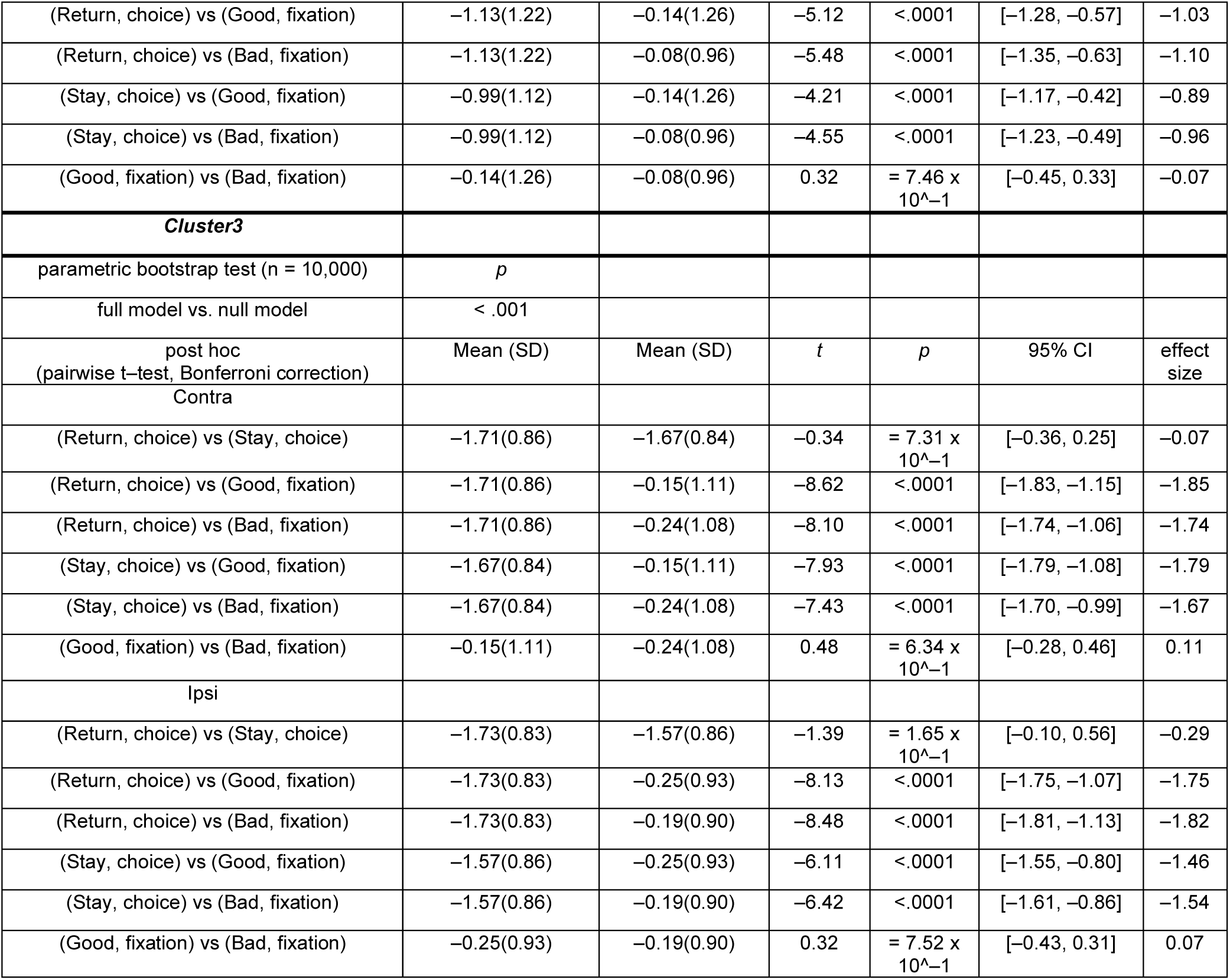
Summary of statistical test to compare the normalized neuronal activity of GPe neurons of clusters 1, 2, and 3 among Return, Stay during choice task, and fixation task in Figure 4.

**Table S10.**
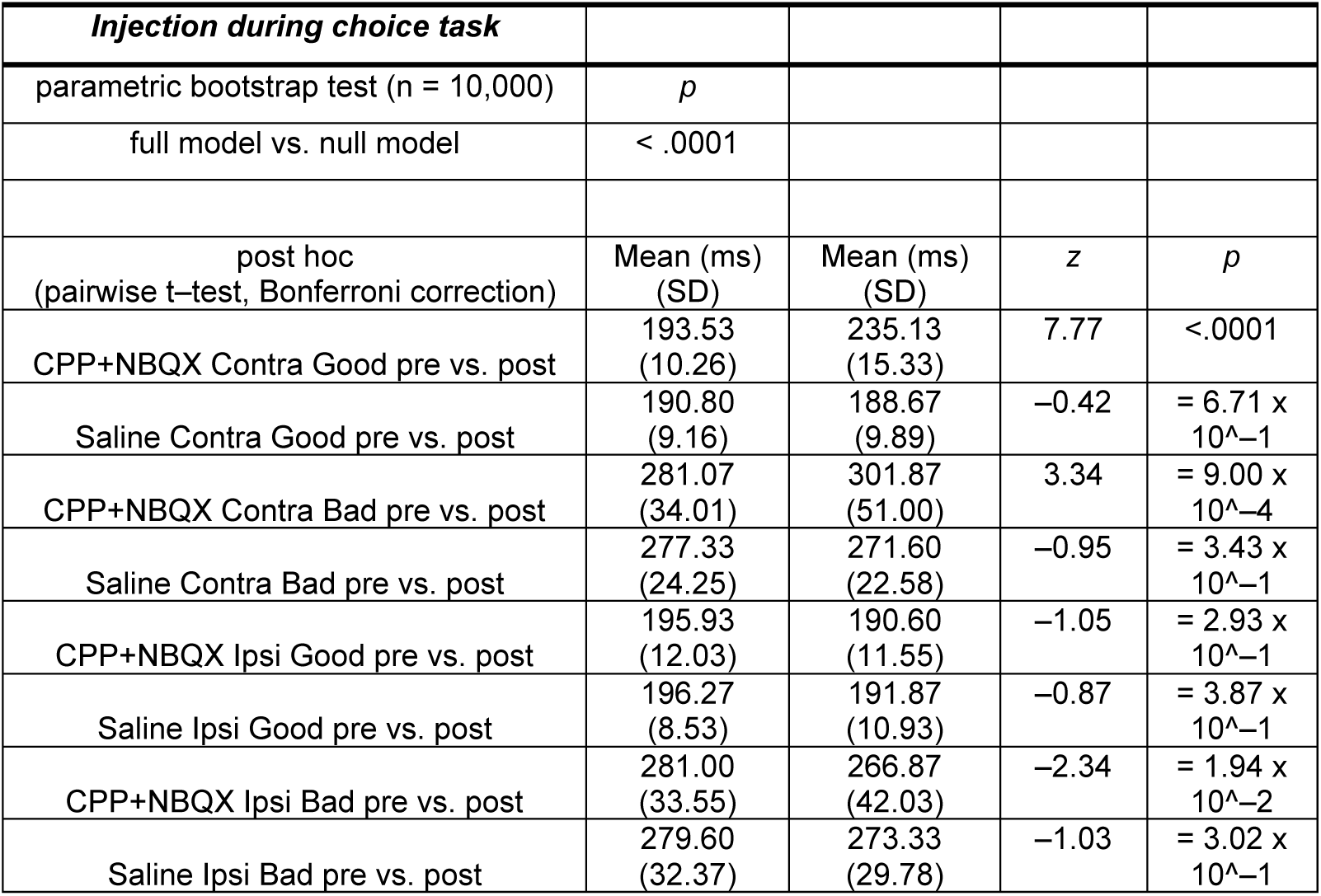
Summary of statistical test to compare the effects of CPP + NBQX injection into GPe during choice task in Figure 5.

**Table S11.**
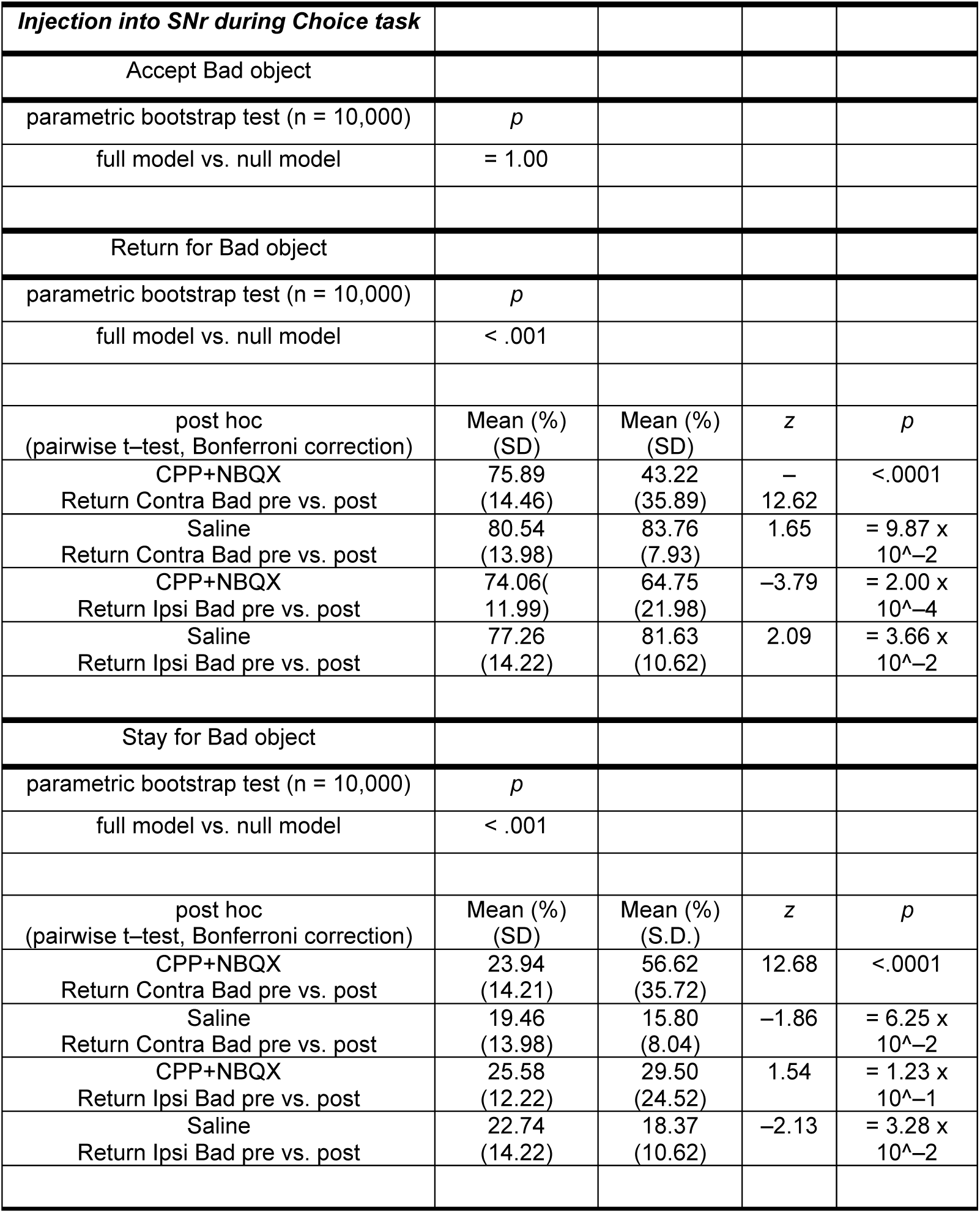
Summary of statistical test to compare the effects of CPP + NBQX injection into GPe while monkeys chose actions for Bad object during Choice task in Figure 5.

## References

1. Graybiel, A.M., Aosaki, T., Flaherty, A. W., & Kimura, M. The basal ganglia and adaptive motor control. Science. 265, 1286–1231 (1994).

2. DeLong, M. R. Primate models of movement disorders of basal ganglia origin. Trends Neurosci. 13, 281–285 (1990).

3. Alexander, G. E. & Crutcher, M. D. Functional architecture of basal ganglia circuits: neural substrates of parallel processing. Trends Neurosci. 13, 266– 271 (1990).

4. Albin, R. L., Young, A. B. & Penney, J. B. The functional anatomy of basal ganglia disorders. Trends Neurosci. 12, 366–375 (1989).

5. Nambu, A. & Chiken, S. External segment of the globus pallidus in health and disease: Its interactions with the striatum and subthalamic nucleus. Neurobiol. Dis. 190, 106362 (2024).

6. Courtney, C. D., Pamukcu, A. & Chan, C. S. Cell and circuit complexity of the external globus pallidus. Nat. Neurosci. 26, 1147–1159 (2023).

7. Fang, L. Z. & Creed, M. C. Updating the striatal-pallidal wiring diagram. Nat. Neurosci. 27, 15–27 (2024).

8. Giossi, C. et al. Rethinking the external globus pallidus and information flow in cortico-basal ganglia-thalamic circuits. Eur. J. Neurosci. 60, 6129–6144 (2024).

9. Mallet, N. et al. Dichotomous organization of the external globus pallidus. Neuron 74, 1075–1086 (2012).

10. Abdi, A. et al. Prototypic and arkypallidal neurons in the dopamine-intact external globus pallidus. J. Neurosci. 35, 6667–6688 (2015).

11. Glajch, K. E. et al. Npas1+ pallidal neurons target striatal projection neurons. J. Neurosci. 36, 5472–5488 (2016).

12. Aristieta, A. et al. A disynaptic circuit in the globus pallidus controls locomotion inhibition. Curr. Biol. 31, 707–721 (2021).

13. Karube, F. et al. Motor cortex can directly drive the globus pallidus neurons in a projection neuron type-dependent manner in the rat. eLife 8, e49511 (2019).

14. Abecassis, Z. A. et al. Npas1+-Nkx2.1+ neurons are an integral part of the cortico-pallido-cortical loop. J. Neurosci. 40, 743–768 (2020).

15. Saunders, A., et al. A direct GABAergic output from the basal ganglia to frontal cortex. Nature 521, 85–89 (2015).

16. Yasukawa, T. et al. Rat intralaminar thalamic nuclei projections to the globus pallidus: a biotinylated dextran amine anterograde tracing study. J. Comp. Neurol. 471, 153–167 (2004).

17. Parent, M. & Parent, A. Single-axon tracing and three-dimensional reconstruction of centre median-parafascicular thalamic neurons in primates. J. Comp. Neurol. 481, 127–144 (2005).

18. DeLong, M. R. Activity of pallidal neurons during movement. J. Neurophysiol. 34, 414–427 (1971).

19. Anderson, M. E. & Horak, F. B. Influence of the globus pallidus on arm movements in monkeys. III. Timing of movement-related information. J. Neurophysiol. 54, 433–448 (1985).

20. Mitchell, S. J., Richardson, R. T., Baker, F. H. & DeLong, M. R. The primate globus pallidus: Neuronal activity related to direction of movement. Exp. Brain Res. 68, 491–505 (1987).

21. Turner, R. S. & Anderson, M. E. Context-dependent modulation of movement-related discharge in the primate globus pallidus. J. Neurosci. 25, 2965–2976 (2005).

22. Saga, Y. et al. Representation of spatial- and object-specific behavioral goals in the dorsal globus pallidus of monkeys during reaching movement. J. Neurosci. 33, 16360–16371 (2013).

23. Goldberg, J. A. & Bergman, H. Computational physiology of the neural networks of the primate globus pallidus: function and dysfunction. Neuroscience. 198, 171–92 (2011).

24. Yoshida, A. & Tanaka, M. Two types of neurons in the primate globus pallidus external segment play distinct roles in antisaccade generation. Cereb. Cortex 26, 1187–1199 (2016).

25. Yoshida, A. & Tanaka, M. Enhanced modulation of neuronal activity during antisaccades in the primate globus pallidus. Cereb. Cortex 19, 206–217 (2009).

26. Aron, A. R. From reactive to proactive and selective control: developing a richer model for stopping inappropriate responses. Biol. Psychiatry 69, e55– e68 (2011).

27. Jahanshahi, M. et al. A fronto-striato-subthalamic-pallidal network for goal-directed and habitual inhibition. Nat. Rev. Neurosci. 16, 719–732 (2015).

28. Dunovan, K. et al. Competing basal ganglia pathways determine the difference between stopping and deciding not to go. eLife 4, e08723 (2015).

29. Gu, B. M. et al. Globus pallidus dynamics reveal covert strategies for behavioral inhibition. eLife 9, e57215 (2020).

30. Yoshida, A. & Hikosaka, О. Involvement of neurons in the non-human primate anterior striatum in proactive inhibition. J. Neurosci. 44, e0866242024 (2024).

31. Shink, E., Bevan, M. D., Bolam, J. P. & Smith, Y. The subthalamic nucleus and the external pallidum: two tightly interconnected structures that control the output of the basal ganglia in the monkey. Neuroscience 69, 745–757 (1996).

32. Nambu, A. et al. Excitatory cortical inputs to pallidal neurons via the subthalamic nucleus in the monkey. J. Neurophysiol. 84, 289–300 (2000).

33. Parent, A. & Hazrati, L. N. Functional anatomy of the basal ganglia. II. The place of subthalamic nucleus and external pallidum in basal ganglia circuitry. Brain Res. Rev. 20, 128–154 (1995).

34. Hunt, A. J. et al. Paraventricular hypothalamic and amygdalar CRF neurons synapse in the external globus pallidus. Brain Struct. Funct. 223, 2685–2698 (2018).

35. Cui, Q. et al. Striatal direct pathway targets Npas1+ pallidal neurons. J. Neurosci. 41, 3966–3987 (2021).

36. Ketzef, M. & Silberberg, G. Differential synaptic input to external globus pallidus neuronal subpopulations in vivo. Neuron 109, 516–529 (2021).

37. Pamukcu, A. et al. Parvalbumin+ and Npas1+ pallidal neurons have distinct circuit topology and function. J. Neurosci. 40, 7855–7876 (2020).

38. Liu, C. et al. Susceptibility-weighted imaging and quantitative susceptibility mapping in the brain. J. Magn. Reson. Imaging 42, 23–41 (2015).

39. Wang, Y. & Liu, T. Quantitative susceptibility mapping (QSM): decoding MRI data for a tissue magnetic biomarker. Magn. Reson. Med. 73, 82–101 (2015).

40. Yoshida, A. et al. Visualization of iron-rich subcortical structures in non-human primates in vivo by quantitative susceptibility mapping at 3T MRI. Neuroimage 241, 118429 (2021)

41. Liu, T. et al. Morphology enabled dipole inversion for quantitative susceptibility mapping using structural consistency between the magnitude image and the susceptibility map. Neuroimage 59, 2560–2568 (2012).

42. Vanegas, M. I. et al. Microinjectrode System for Combined Drug Infusion and Electrophysiology. J. Vis. Exp. 153, e60365 (2019).

43. Tachibana, Y. et al. Subthalamo-pallidal interactions underlying parkinsonian neuronal oscillations in the primate basal ganglia. Eur. J. Neurosci. 34, 1470– 1484 (2011).

44. Adler, A. et al. Temporal convergence of dynamic cell assemblies in the striato-pallidal network. J. Neurosci. 32, 2473–2484 (2012).

45. Kaplan, A. D. et al. Dissociable roles of ventral pallidum neurons in the basal ganglia reinforcement learning network. Nat. Neurosci. 23, 556–564 (2020).

46. Yu, Z. et al. Beyond t test and ANOVA: applications of mixed-effects models for more rigorous statistical analysis in neuroscience research. Neuron 110, 21–35 (2022).

47. Bates, D., Maechler, B., Bolker, B. & Walker, S. F. Fitting linear mixed-effects models using lme4. J. Stat. Softw. 67, 1–48 (2014).

48. Halekoh, U. & Højsgaard, S. A Kenward-Roger approximation and parametric bootstrap methods for tests in linear mixed models – the R package pbkrtest. J. Stat. Softw. 59, 1–30 (2014).

49. Lenth, R. et al. Estimated marginal means, aka least-squares means. R package version 1.3.2 (2019).

50. Bürkner, P.-C. brms: An R Package for Bayesian Multilevel Models Using Stan. J. Stat. Softw. 80, 1–28 (2017).

