## Supplemental materials for "Excitatory drive to the globus pallidus external segment facilitates action initiation in non-human primates"

Multiple comparisons:
8 pairwise comparisons (pre vs. post for each condition)
Bonferroni-corrected threshold: $\alpha = 0.05/8$

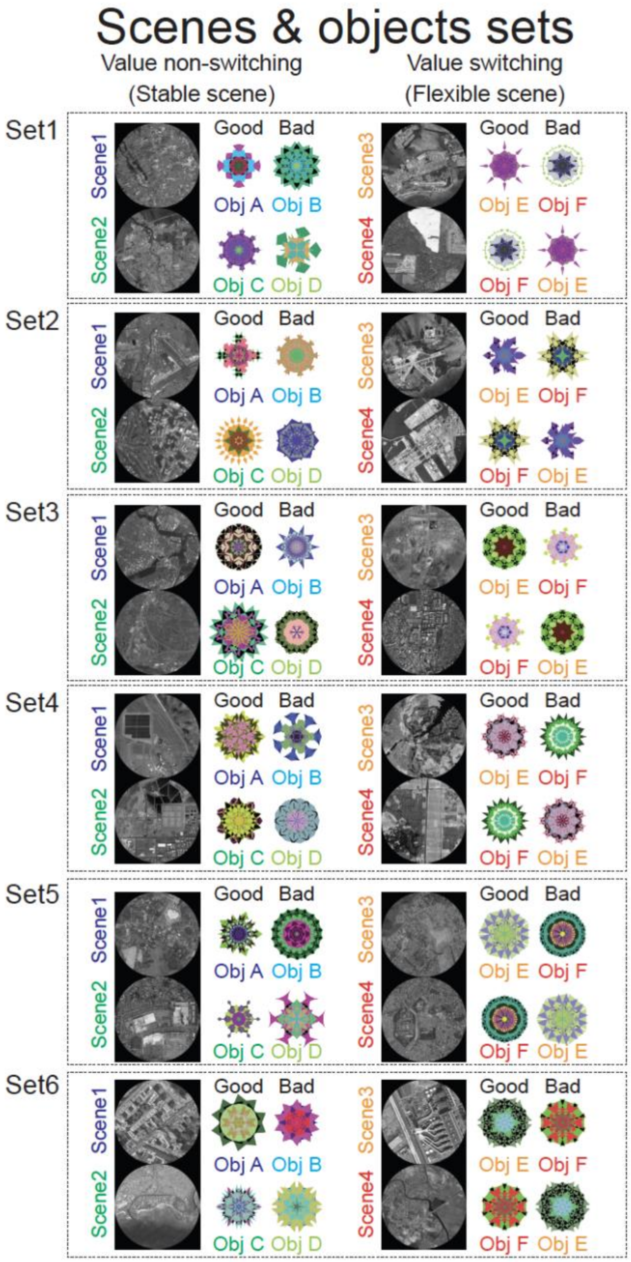

**Figure S1. All sets of scenes 1-4 and good and bad objects for the choice task.**

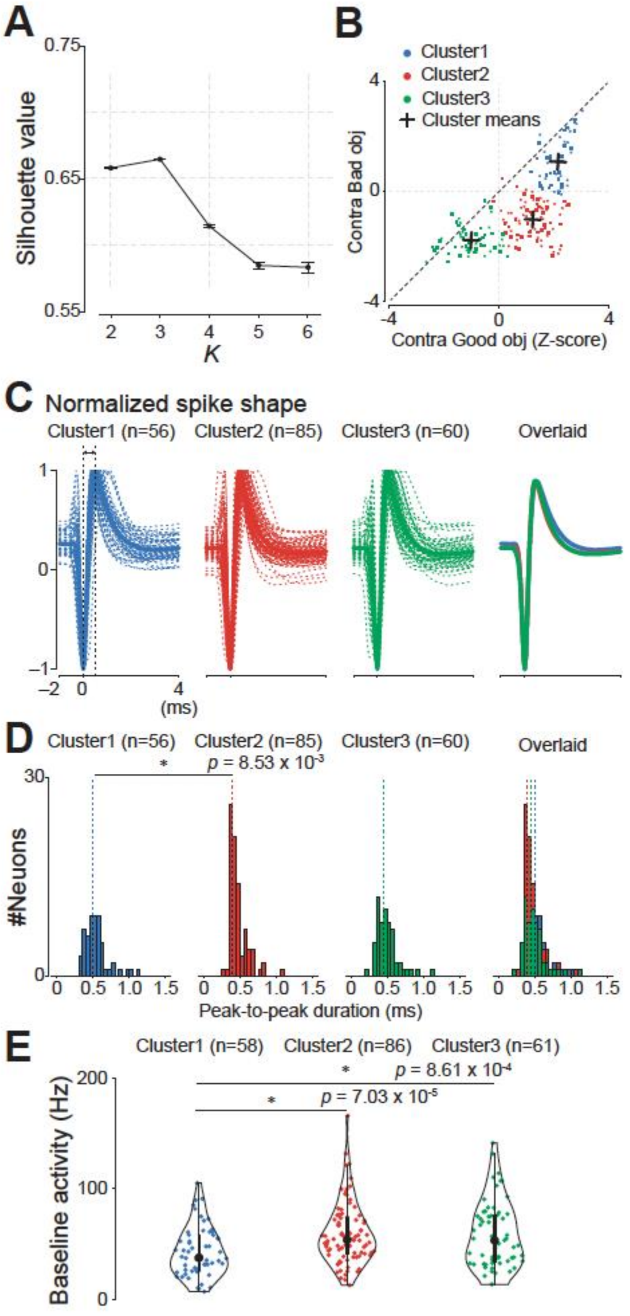

**Figure S2. Cluster analysis of GPe neurons based on task-related activity and comparison of their electrophysiological properties.**

(A) Determination of the optimal number of clusters (K) for k-means clustering. The graph shows the mean ± SD silhouette values for different K values, calculated from 5,000 simulations. The silhouette value was significantly higher for K=3 than for other values ($p < 0.001$, one-way ANOVA), indicating that the neurons were best classified into three clusters.

(B) Scatter plot of the average standardized firing rates of each neuron during the presentation of a contralateral good object (x-axis) and a contralateral bad object (y-axis). The firing rates were calculated for the period between 100 and 300 ms after object presentation and standardized using the z-score method (see Methods for details). Each dot represents a neuron, and the colors indicate the cluster to which the neuron was assigned: Cluster 1 (blue), Cluster 2 (red), and Cluster 3 (green). The crosses indicate the mean values for each cluster.

(C) Normalized spike shapes for each cluster. The thick lines represent the average spike shape for each cluster, and the thin lines represent the individual spike shapes. The spike shapes were normalized to the trough and peak amplitudes for each neuron.

(D) Histograms of the peak-to-peak duration for each cluster. The peak-to-peak duration was defined as the time between the trough and the peak of the spike waveform. The asterisk indicates a significant difference between Clusters 1 and 2 ($p = 8.53 \times 10^{-3}$, Kruskal–Wallis test followed by Dunn's post-hoc test).

(E) Violin plots of the baseline firing rates for each cluster. The baseline firing rate was defined as the average firing rate during the 500 ms period preceding the scene onset. The format of the violin plots is the same as that in Figure 2B. The larger circle indicates the median value, the thick vertical line shows the interquartile range (IQR), and the thin vertical line indicates the range from the lower to the upper adjacent values (1.5xIQR below the first quartile and 1.5xIQR above the third quartile, respectively). The asterisks indicate significant differences between Clusters 1 and 2 ($p = 7.03 \times 10^{-5}$) and between Clusters 1 and 3 ($p = 8.61 \times 10^{-4}$) (Kruskal–Wallis test followed by Dunn's post-hoc test).

Abbreviations: GPe, external segment of the globus pallidus; SD, standard deviation.

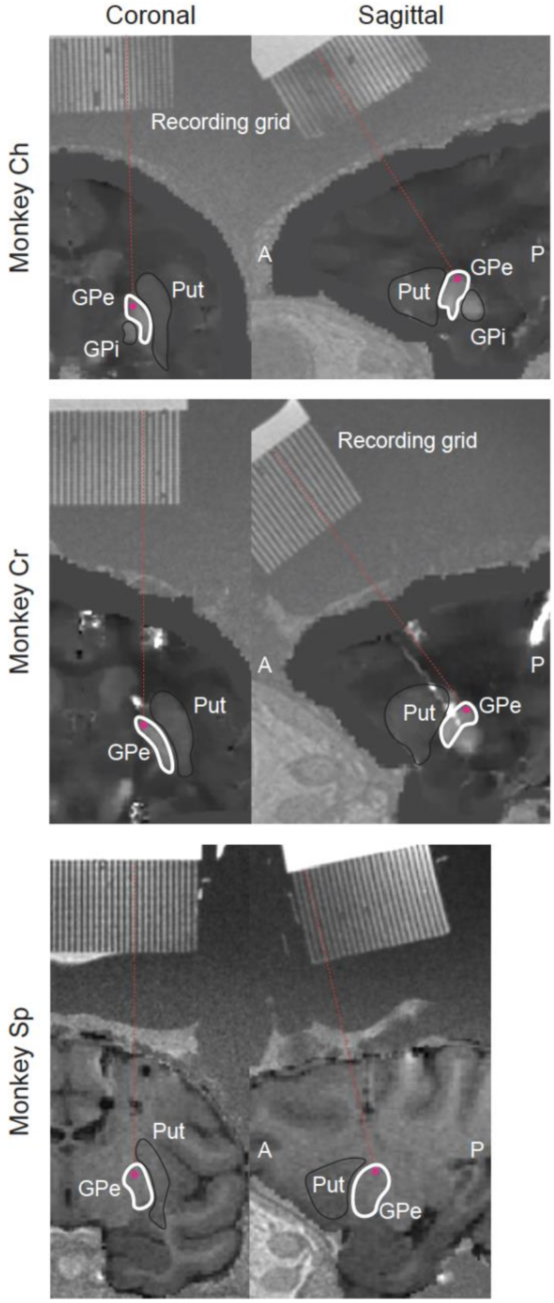

**Figure S3. Examples of injection sites in the GPe of three macaque monkeys (Monkey Ch, Monkey Cr, and Monkey Sp).**
Coronal (left) and sagittal (right) sections of MRI images show representative injection sites (red dots) in each monkey. The MRI images were fused images of

high-resolution T1-weighted images and either QSM or a conventional three-dimensional T1-weighted image. The white lines delineate the approximate borders of the GPe and putamen. The red dots indicate the location of the injection site, which was targeted to be in the cdGPe. Abbreviations: A, anterior; P, posterior; cdGPe, caudodorsal globus pallidus external segment; GPe, globus pallidus external segment; GPi, globus pallidus internal segment; Put, putamen; QSM, quantitative susceptibility mapping; MRI, magnetic resonance imaging.

**Table S1. Saccade reaction times in each condition of each monkey.**

| **Monkey Cr** | n | mean RT (ms) | SD | 95% CI |
|---|---|---|---|---|
| **Good obj** | | | | |
| ObjA in Scene1 | 2176 | 170.5 | 23.7 | [169.5 171.5] |
| ObjC in Scene2 | 2225 | 171.6 | 24.2 | [170.5 172.6] |
| ObjE in Scene3 | 2188 | 180.0 | 24.4 | [179.0 181.0] |
| ObjF in Scene4 | 2163 | 174.4 | 22.6 | [173.4 175.3] |
| **Bad obj** | | | | |
| ObjB in Scene1 | 1699 | 250.0 | 50.4 | [247.6 252.4] |
| ObjD in Scene2 | 1769 | 254.1 | 48.9 | [251.8 256.4] |
| ObjF in Scene3 | 2003 | 232.3 | 51.6 | [230.1 234.6] |
| ObjE in Scene4 | 1958 | 235.7 | 55.2 | [233.2 238.1] |
| **Monkey Sp** | n | mean RT (ms) | SD | 95% CI |
| **Good obj** | | | | |
| ObjA in Scene1 | 1960 | 168.8 | 24.2 | [167.8 169.9] |
| ObjC in Scene2 | 1947 | 174.9 | 25.9 | [173.8 176.2] |
| ObjE in Scene3 | 1934 | 179.0 | 24.0 | [177.9 180.1] |
| ObjF in Scene4 | 1932 | 178.4 | 25.6 | [177.3 179.6] |
| **Bad obj** | | | | |
| ObjB in Scene1 | 1385 | 284.1 | 45.5 | [281.7 286.5] |
| ObjD in Scene2 | 1270 | 291.4 | 42.4 | [289.0 293.7] |
| ObjF in Scene3 | 1629 | 267.7 | 54.3 | [265.1 270.3] |
| ObjE in Scene4 | 1452 | 280.3 | 52.1 | [277.6 283.0] |

**Table S2. Counts of chosen actions for Bad objects**

| **Monkey Cr** | total | accept | Return | Stay | other | fxbreak |
|---|---|---|---|---|---|---|
| Scene1 | 2219 | 10 | 1727 | 437 | 3 | 42 |
| Scene2 | 2206 | 4 | 1791 | 398 | 0 | 13 |
| Scene3 | 2150 | 7 | 2009 | 119 | 1 | 14 |
| Scene4 | 2255 | 5 | 1973 | 239 | 0 | 38 |
| non-switch(scene1,2) | 4425 | 14 | 3518 | 835 | 4 | 55 |
| switch(scene3,4) | 4405 | 12 | 3982 | 358 | 1 | 52 |
| **Monkey Sp** | total | accept | Return | Stay | other | fxbreak |
| Scene1 | 1927 | 6 | 1357 | 556 | 0 | 8 |
| Scene2 | 1892 | 9 | 1246 | 629 | 0 | 8 |
| Scene3 | 2032 | 3 | 1616 | 407 | 0 | 6 |
| Scene4 | 1816 | 4 | 1435 | 365 | 0 | 12 |
| non-switch(scene1,2) | 3819 | 15 | 2603 | 1185 | 0 | 16 |
| switch(scene3,4) | 3848 | 7 | 3051 | 772 | 0 | 18 |

**Table S3. Summary of statistical test to compare the normalized neuronal activity of GPe neurons of cluster1 at target onset among conditions during choice task in Figure 2.**

| ***Cluster1*** | | | | | | |
|---|---|---|---|---|---|---|
| parametric bootstrap test (n = 10,000) | *p* | | | | | |
| full model vs. null model | < .001 | | | | | |
| post hoc (pairwise t–test, Bonferroni correction) | Mean (SD) | Mean (SD) | *t* | *p* | 95% CI | effect size |
| **Scene1** | | | | | | |
| (good, contra) vs (bad, contra) | 2.17(0.44) | 1.01(1.17) | 8.42 | <.0001 | [0.90, 1.44] | 1.56 |
| (good, contra) vs (good, ipsi) | 2.17(0.44) | 1.20(0.86) | 7.00 | <.0001 | [0.70, 1.24] | 1.30 |
| (good, contra) vs (bad, ipsi) | 2.17(0.44) | 0.41(1.35) | 12.71 | <.0001 | [1.49, 2.03] | 2.36 |
| (bad, contra) vs (good, ipsi) | 1.01(1.17) | 1.20(0.86) | –1.42 | = 1.55 x 10^–1 | [–0.47, 0.07] | –0.26 |
| (bad, contra) vs (bad, ipsi) | 1.01(1.17) | 0.41(1.35) | 4.29 | <.0001 | [0.32, 0.87] | 0.80 |
| (good, ipsi) vs (bad, ipsi) | 1.20(0.86) | 0.41(1.35) | 5.71 | <.0001 | [0.52, 1.06] | 1.06 |
| **Scene2** | | | | | | |
| (good, contra) vs (bad, contra) | 2.23(0.47) | 0.95(1.08) | 9.23 | <.0001 | [1.01, 1.55] | 1.71 |
| (good, contra) vs (good, ipsi) | 2.23(0.47) | 1.46(0.75) | 5.51 | <.0001 | [0.49, 1.04] | 1.02 |
| (good, contra) vs (bad, ipsi) | 2.23(0.47) | 0.50(1.07) | 12.47 | <.0001 | [1.46, 2.00] | 2.32 |
| (bad, contra) vs (good, ipsi) | 0.95(1.08) | 1.46(0.75) | –3.73 | = 2.00 x 10^–4 | [–0.79, –0.24] | –0.69 |
| (bad, contra) vs (bad, ipsi) | 0.95(1.08) | 0.50(1.07) | 3.24 | 1.20 x 10^–3 | [0.18, 0.72] | 0.60 |
| (good, ipsi) vs (bad, ipsi) | 1.46(0.75) | 0.50(1.07) | 6.96 | <.0001 | [0.69, 1.24] | 1.29 |
| **Scene3** | | | | | | |
| (good, contra) vs (bad, contra) | 2.14(0.59) | 1.29(1.07) | 6.12 | <.0001 | [0.58, 1.12] | 1.14 |
| (good, contra) vs (good, ipsi) | 2.14(0.59) | 1.24(0.96) | 6.50 | <.0001 | [0.63, 1.17] | 1.21 |
| (good, contra) vs (bad, ipsi) | 2.14(0.59) | 0.58(1.24) | 11.24 | <.0001 | [1.29, 1.83] | 2.09 |
| (bad, contra) vs (good, ipsi) | 1.29(1.07) | 1.24(0.96) | 0.38 | = 7.05 x 10^–1 | [–0.22, 0.32] | 0.07 |
| (bad, contra) vs (bad, ipsi) | 1.29(1.07) | 0.58(1.24) | 5.12 | <.0001 | [0.44, 0.98] | 0.95 |
| (good, ipsi) vs (bad, ipsi) | 1.24(0.96) | 0.58(1.24) | 4.74 | <.0001 | [0.39, 0.93] | 0.88 |
| **Scene4** | | | | | | |
| (good, contra) vs (bad, contra) | 2.15(0.55) | 1.17(1.06) | 7.05 | <.0001 | [0.71, 1.25] | 1.31 |
| (good, contra) vs (good, ipsi) | 2.15(0.55) | 1.23(0.98) | 6.59 | <.0001 | [0.64, 1.19] | 1.22 |
| (good, contra) vs (bad, ipsi) | 2.15(0.55) | 0.51(1.08) | 11.83 | <.0001 | [1.37, 1.91] | 2.20 |
| (bad, contra) vs (good, ipsi) | 1.17(1.06) | 1.23(0.98) | 0.46 | = 6.45 x 10^–1 | [–0.34, 0.21] | –0.09 |
| (bad, contra) vs (bad, ipsi) | 1.17(1.06) | 0.51(1.08) | 4.78 | <.0001 | [0.39, 0.94] | 0.89 |
| (good, ipsi) vs (bad, ipsi) | 1.23(0.98) | 0.51(1.08) | 5.24 | <.0001 | [0.46, 1.00] | 0.97 |

**Table S4. Summary of statistical test to compare the normalized neuronal activity of GPe neurons of cluster2 at target onset among conditions during choice task in Figure 2.**

| ***Cluster2*** | | | | | | |
|---|---|---|---|---|---|---|
| parametric bootstrap test (n = 10,000) | *p* | | | | | |
| full model vs. null model | < .001 | | | | | |
| post hoc (pairwise t–test, Bonferroni correction) | Mean (SD) | Mean (SD) | *t* | *p* | 95% CI | effect size |
| **Scene1** | | | | | | |
| (good, contra) vs (bad, contra) | 1.33(0.68) | –1.11(1.08) | 16.50 | <.0001 | [2.14, 2.72] | 2.52 |
| (good, contra) vs (good, ipsi) | 1.33(0.68) | 0.34(0.94) | 6.72 | <.0001 | [0.70, 1.28] | 1.02 |
| (good, contra) vs (bad, ipsi) | 1.33(0.68) | –1.13(1.22) | 16.67 | <.0001 | [2.17, 2.74] | 2.54 |
| (bad, contra) vs (good, ipsi) | –1.11(1.08) | 0.34(0.94) | –9.79 | <.0001 | [–1.73, –1.15] | –1.49 |
| (bad, contra) vs (bad, ipsi) | –1.11(1.08) | –1.13(1.22) | 0.16 | = 8.70 x 10^–1 | [–0.27, 0.32] | 0.02 |
| (good, ipsi) vs (bad, ipsi) | 0.34(0.94) | –1.13(1.22) | 9.95 | <.0001 | [1.18, 1.76] | 1.52 |
| **Scene2** | | | | | | |
| (good, contra) vs (bad, contra) | 1.22(0.82) | –1.09(1.01) | 15.73 | <.0001 | [2.03, 2.61] | 2.40 |
| (good, contra) vs (good, ipsi) | 1.22(0.82) | 0.39(1.12) | 5.67 | <.0001 | [0.55, 1.12] | 0.86 |
| (good, contra) vs (bad, ipsi) | 1.22(0.82) | –1.12(1.05) | 15.95 | <.0001 | [2.06, 2.64] | 2.43 |
| (bad, contra) vs (good, ipsi) | –1.09(1.01) | 0.39(1.12) | –10.06 | <.0001 | [–1.77, –1.19] | –1.53 |
| (bad, contra) vs (bad, ipsi) | –1.09(1.01) | –1.12(1.05) | 0.22 | = 8.29 x 10^–1 | [–0.26, 0.32] | 0.03 |
| (good, ipsi) vs (bad, ipsi) | 0.39(1.12) | –1.12(1.05) | 10.28 | <.0001 | [1.23, 1.80] | 1.57 |
| **Scene3** | | | | | | |
| (good, contra) vs (bad, contra) | 1.31(0.83) | –0.90(1.13) | 15.02 | <.0001 | [1.92, 2.50] | 2.29 |
| (good, contra) vs (good, ipsi) | 1.31(0.83) | 0.52(1.12) | 5.43 | <.0001 | [0.51, 1.09] | 0.83 |
| (good, contra) vs (bad, ipsi) | 1.31(0.83) | –1.05(1.11) | 16.07 | <.0001 | [2.08, 2.66] | 2.45 |
| (bad, contra) vs (good, ipsi) | –0.90(1.13) | 0.52(1.12) | –9.59 | <.0001 | [–1.70, –1.12] | –1.46 |
| (bad, contra) vs (bad, ipsi) | –0.90(1.13) | –1.05(1.11) | 1.05 | = 2.94 x 10^–1 | [–0.13, 0.44] | 0.16 |
| (good, ipsi) vs (bad, ipsi) | 0.52(1.12) | –1.05(1.11) | 10.64 | <.0001 | [1.28, 1.86] | 1.62 |
| **Scene4** | | | | | | |
| (good, contra) vs (bad, contra) | 1.17(0.88) | –1.05(1.26) | 15.04 | <.0001 | [1.93, 2.50] | 2.29 |
| (good, contra) vs (good, ipsi) | 1.17(0.88) | 0.39(1.10) | 5.26 | <.0001 | [0.49, 1.06] | 0.80 |
| (good, contra) vs (bad, ipsi) | 1.17(0.88) | –1.19(1.18) | 15.98 | <.0001 | [2.06, 2.64] | 2.44 |
| (bad, contra) vs (good, ipsi) | –1.05(1.26) | 0.39(1.10) | –9.77 | <.0001 | [–1.73, –1.15] | –1.49 |
| (bad, contra) vs (bad, ipsi) | –1.05(1.26) | –1.19(1.18) | 0.94 | = 3.46 x 10^–1 | [–0.15, 0.43] | 0.14 |
| (good, ipsi) vs (bad, ipsi) | 0.39(1.10) | –1.19(1.18) | 10.72 | <.0001 | [1.29, 1.87] | 1.63 |

**Table S5. Summary of statistical test to compare the normalized neuronal activity of GPe neurons of cluster3 at target onset among conditions during choice task in Figure 2.**

| ***Cluster3*** | | | | | | |
|---|---|---|---|---|---|---|
| parametric bootstrap test (n = 10,000) | *p* | | | | | |
| full model vs. null model | < .001 | | | | | |
| post hoc (pairwise t–test, Bonferroni correction) | Mean (SD) | Mean (SD) | *t* | *p* | 95% CI | effect size |
| **Scene1** | | | | | | |
| (good, contra) vs (bad, contra) | –0.93(1.01) | –1.70(0.86) | 6.00 | <.0001 | [0.52, 1.02] | 1.09 |
| (good, contra) vs (good, ipsi) | –0.93(1.01) | –1.14(1.07) | 1.59 | = 1.13 x 10^–1 | [–0.05, 0.46] | 0.29 |
| (good, contra) vs (bad, ipsi) | –0.93(1.01) | –1.72(0.83) | 6.15 | <.0001 | [0.54, 1.04] | 1.11 |
| (bad, contra) vs (good, ipsi) | –1.70(0.86) | –1.14(1.07) | –4.41 | <.0001 | [–0.82, –0.32] | –0.80 |
| (bad, contra) vs (bad, ipsi) | –1.70(0.86) | –1.72(0.83) | 0.15 | = 8.78 x 10^–1 | [–0.23, 0.27] | 0.03 |
| (good, ipsi) vs (bad, ipsi) | –1.14(1.07) | –1.72(0.83) | 4.57 | <.0001 | [0.33, 0.84] | 0.83 |
| **Scene2** | | | | | | |
| (good, contra) vs (bad, contra) | –1.14(0.96) | –1.72(0.81) | 4.59 | <.0001 | [0.34, 0.84] | 0.83 |
| (good, contra) vs (good, ipsi) | –1.14(0.96) | –1.30(1.02) | 1.27 | = 2.05 x 10^–1 | [–0.09, 0.42] | 0.23 |
| (good, contra) vs (bad, ipsi) | –1.14(0.96) | –1.87(0.83) | 5.61 | <.0001 | [0.47, 0.98] | 1.02 |
| (bad, contra) vs (good, ipsi) | –1.72(0.81) | –1.30(1.02) | –3.32 | = 9.00 x 10^–4 | [–0.68, –0.17] | –0.60 |
| (bad, contra) vs (bad, ipsi) | –1.72(0.81) | –1.87(0.83) | 1.05 | = 2.96 x 10^–1 | [–0.12, 0.39] | 0.19 |
| (good, ipsi) vs (bad, ipsi) | –1.30(1.02) | –1.87(0.83) | 4.35 | <.0001 | [0.31, 0.81] | 0.79 |
| **Scene3** | | | | | | |
| (good, contra) vs (bad, contra) | –1.06(0.89) | –1.78(0.87) | 5.60 | <.0001 | [0.47, 0.97] | 1.01 |
| (good, contra) vs (good, ipsi) | –1.06(0.89) | –1.33(0.99) | 2.08 | = 3.82 x 10^–2 | [0.01, 0.52] | 0.38 |
| (good, contra) vs (bad, ipsi) | –1.06(0.89) | –1.76(0.90) | 5.45 | <.0001 | [0.45, 0.95] | 0.99 |
| (bad, contra) vs (good, ipsi) | –1.78(0.87) | –1.33(0.99) | –3.52 | = 5.00 x 10^–4 | [–0.70, –0.20] | –0.64 |
| (bad, contra) vs (bad, ipsi) | –1.78(0.87) | –1.76(0.90) | –0.15 | = 8.85 x 10^–1 | [–0.27, 0.23] | –0.03 |
| (good, ipsi) vs (bad, ipsi) | –1.33(0.99) | –1.76(0.90) | 3.38 | = 8.00 x 10^–4 | [0.18, 0.69] | 0.61 |
| **Scene4** | | | | | | |
| (good, contra) vs (bad, contra) | –1.00(1.01) | –1.83(0.94) | 6.47 | <.0001 | [0.58, 1.08] | 1.17 |
| (good, contra) vs (good, ipsi) | –1.00(1.01) | –1.20(1.01) | 1.55 | = 1.21 x 10^–1 | [–0.05, 0.45] | 0.28 |
| (good, contra) vs (bad, ipsi) | –1.00(1.01) | –1.78(0.94) | 6.07 | <.0001 | [0.53, 1.03] | 1.10 |
| (bad, contra) vs (good, ipsi) | –1.83(0.94) | –1.20(1.01) | –4.91 | <.0001 | [–0.88, –0.38] | –0.89 |
| (bad, contra) vs (bad, ipsi) | –1.83(0.94) | –1.78(0.94) | –0.39 | = 6.95 x 10^–1 | [–0.30, 0.20] | –0.07 |
| (good, ipsi) vs (bad, ipsi) | –1.20(1.01) | –1.78(0.94) | 4.52 | <.0001 | [0.33, 0.83] | 0.82 |

**Table S6. Summary of statistical test to compare the normalized neuronal activity of GPe neurons of cluster1 at saccade onset among conditions during choice task in Figure 3.**

| ***Cluster1*** | | | | | | |
|---|---|---|---|---|---|---|
| parametric bootstrap test (n = 10,000) | *p* | | | | | |
| full model vs. null model | < .001 | | | | | |
| post hoc (pairwise t–test, Bonferroni correction) | Mean (SD) | Mean (SD) | *t* | *p* | 95% CI | effect size |
| **Scene1** | | | | | | |
| (good, contra) vs (bad, contra) | 1.72(0.58) | 0.85(1.19) | 5.88 | <.0001 | [0.58, 1.16] | 1.09 |
| (good, contra) vs (good, ipsi) | 1.72(0.58) | 0.81(0.88) | 6.15 | <.0001 | [0.62, 1.20] | 1.14 |
| (good, contra) vs (bad, ipsi) | 1.72(0.58) | 0.35(1.35) | 9.22 | <.0001 | [1.08, 1.66] | 1.71 |
| (bad, contra) vs (good, ipsi) | 0.85(1.19) | 0.81(0.88) | 0.27 | = 7.86 x 10^–1 | [–0.25, 0.33] | 0.05 |
| (bad, contra) vs (bad, ipsi) | 0.85(1.19) | 0.35(1.35) | 3.34 | = 9.00 x 10^–4 | [0.21, 0.79] | 0.62 |
| (good, ipsi) vs (bad, ipsi) | 0.81(0.88) | 0.35(1.35) | 3.07 | = 2.20 x 10^–3 | [0.16, 0.75] | 0.57 |
| **Scene2** | | | | | | |
| (good, contra) vs (bad, contra) | 1.77(0.55) | 0.82(1.17) | 6.38 | <.0001 | [0.66, 1.24] | 1.18 |
| (good, contra) vs (good, ipsi) | 1.77(0.55) | 1.16(0.90) | 4.09 | <.0001 | [0.32, 0.90] | 0.76 |
| (good, contra) vs (bad, ipsi) | 1.77(0.55) | 0.46(1.08) | 8.78 | <.0001 | [1.01, 1.59] | 1.63 |
| (bad, contra) vs (good, ipsi) | 0.82(1.17) | 1.16(0.90) | –2.29 | = 2.20 x 10^–2 | [–0.63, –0.05] | –0.43 |
| (bad, contra) vs (bad, ipsi) | 0.82(1.17) | 0.46(1.08) | 2.40 | = 1.66 x 10^–2 | [0.06, 0.65] | 0.45 |
| (good, ipsi) vs (bad, ipsi) | 1.16(0.90) | 0.46(1.08) | 4.69 | <.0001 | [0.41, 0.99] | 0.87 |
| **Scene3** | | | | | | |
| (good, contra) vs (bad, contra) | 1.74(0.56) | 1.14(1.17) | 4.07 | = 1.00 x 10^–4 | [0.31, 0.90] | 0.76 |
| (good, contra) vs (good, ipsi) | 1.74(0.56) | 0.96(1.00) | 5.25 | <.0001 | [0.49, 1.07] | 0.97 |
| (good, contra) vs (bad, ipsi) | 1.74(0.56) | 0.54(1.23) | 8.08 | <.0001 | [0.91, 1.49] | 1.50 |
| (bad, contra) vs (good, ipsi) | 1.14(1.17) | 0.96(1.00) | 1.18 | = 2.39 x 10^–1 | [–0.12, 0.47] | 0.22 |
| (bad, contra) vs (bad, ipsi) | 1.14(1.17) | 0.54(1.23) | 4.01 | = 1.00 x 10^–4 | [0.30, 0.89] | 0.74 |
| (good, ipsi) vs (bad, ipsi) | 0.96(1.00) | 0.54(1.23) | 2.83 | = 4.70 x 10^–3 | [0.13, 0.71] | 0.53 |
| **Scene4** | | | | | | |
| (good, contra) vs (bad, contra) | 1.69(0.63) | 1.13(1.07) | 3.80 | = 2.00 x 10^–4 | [0.27, 0.86] | 0.71 |
| (good, contra) vs (good, ipsi) | 1.69(0.63) | 0.95(1.01) | 5.05 | <.0001 | [0.46, 1.04] | 0.94 |
| (good, contra) vs (bad, ipsi) | 1.69(0.63) | 0.44(1.17) | 8.48 | <.0001 | [0.97, 1.55] | 1.57 |
| (bad, contra) vs (good, ipsi) | 1.13(1.07) | 0.95(1.01) | 1.25 | = 2.12 x 10^–1 | [–0.11, 0.48] | 0.23 |
| (bad, contra) vs (bad, ipsi) | 1.13(1.07) | 0.44(1.17) | 4.68 | <.0001 | [0.40, 0.99] | 0.87 |
| (good, ipsi) vs (bad, ipsi) | 0.95(1.01) | 0.44(1.17) | 3.43 | = 6.00 x 10^–4 | [0.22, 0.80] | 0.64 |

**Table S7. Summary of statistical test to compare the normalized neuronal activity of GPe neurons of cluster2 at saccade onset among conditions during choice task in Figure 3.**

| ***Cluster2*** | | | | | | |
|---|---|---|---|---|---|---|
| parametric bootstrap test (n = 10,000) | *p* | | | | | |
| full model vs. null model | < .001 | | | | | |
| post hoc<br>(pairwise t–test, Bonferroni correction) | Mean (SD) | Mean (SD) | *t* | *p* | 95% CI | effect size |
| **Scene1** | | | | | | |
| (good, contra) vs (bad, contra) | 0.98(0.73) | –1.21(1.06) | 14.36 | <.0001 | [1.89, 2.48] | 2.19 |
| (good, contra) vs (good, ipsi) | 0.98(0.73) | 0.15(0.99) | 5.42 | <.0001 | [0.53, 1.12] | 0.83 |
| (good, contra) vs (bad, ipsi) | 0.98(0.73) | –1.18(1.23) | 14.22 | <.0001 | [1.86, 2.46] | 2.17 |
| (bad, contra) vs (good, ipsi) | –1.21(1.06) | 0.15(0.99) | –8.94 | <.0001 | [–1.66, –1.06] | –1.36 |
| (bad, contra) vs (bad, ipsi) | –1.21(1.06) | –1.18(1.23) | –0.14 | = 8.86 x 10^–1 | [–0.32, 0.28] | –0.02 |
| (good, ipsi) vs (bad, ipsi) | 0.15(0.99) | –1.18(1.23) | 8.80 | <.0001 | [1.04, 1.64] | 1.34 |
| **Scene2** | | | | | | |
| (good, contra) vs (bad, contra) | 1.02(0.82) | –1.18(1.00) | 14.48 | <.0001 | [1.90, 2.50] | 2.21 |
| (good, contra) vs (good, ipsi) | 1.02(0.82) | 0.33(1.10) | 4.60 | <.0001 | [0.40, 1.00] | 0.70 |
| (good, contra) vs (bad, ipsi) | 1.02(0.82) | –1.17(1.06) | 14.45 | <.0001 | [1.90, 2.49] | 2.20 |
| (bad, contra) vs (good, ipsi) | –1.18(1.00) | 0.33(1.10) | –9.88 | <.0001 | [–1.80, –1.20] | –1.50 |
| (bad, contra) vs (bad, ipsi) | –1.18(1.00) | –1.17(1.06) | –0.03 | = 9.73 x 10^–1 | [–0.30, 0.29] | –0.01 |
| (good, ipsi) vs (bad, ipsi) | 0.33(1.10) | –1.17(1.06) | 9.84 | <.0001 | [1.20, 1.80] | 1.50 |
| **Scene3** | | | | | | |
| (good, contra) vs (bad, contra) | 1.09(0.79) | –0.90(1.14) | 13.13 | <.0001 | [1.70, 2.29] | 2.00 |
| (good, contra) vs (good, ipsi) | 1.09(0.79) | 0.38(1.08) | 4.67 | <.0001 | [0.41, 1.01] | 0.71 |
| (good, contra) vs (bad, ipsi) | 1.09(0.79) | –1.00(1.16) | 13.75 | <.0001 | [1.79, 2.39] | 2.10 |
| (bad, contra) vs (good, ipsi) | –0.90(1.14) | 0.38(1.08) | –8.45 | <.0001 | [–1.58, –0.99] | –1.29 |
| (bad, contra) vs (bad, ipsi) | –0.90(1.14) | –1.00(1.16) | 0.63 | = 5.31 x 10^–1 | [–0.20, 0.39] | 0.10 |
| (good, ipsi) vs (bad, ipsi) | 0.38(1.08) | –1.00(1.16) | 9.08 | <.0001 | [1.08, 1.68] | 1.38 |
| **Scene4** | | | | | | |
| (good, contra) vs (bad, contra) | 0.87(0.86) | –1.11(1.54) | 13.03 | <.0001 | [1.68, 2.28] | 1.99 |
| (good, contra) vs (good, ipsi) | 0.87(0.86) | 0.23(1.04) | 4.21 | <.0001 | [0.34, 0.94] | 0.64 |
| (good, contra) vs (bad, ipsi) | 0.87(0.86) | –1.22(1.27) | 13.76 | <.0001 | [1.79, 2.39] | 2.10 |
| (bad, contra) vs (good, ipsi) | –1.11(1.54) | 0.23(1.04) | –8.82 | <.0001 | [–1.64, –1.04] | –1.34 |
| (bad, contra) vs (bad, ipsi) | –1.11(1.54) | –1.22(1.27) | 0.73 | = 4.66 x 10^–1 | [–0.19, 0.41] | 0.11 |
| (good, ipsi) vs (bad, ipsi) | 0.23(1.04) | –1.22(1.27) | 9.55 | <.0001 | [1.15, 1.75] | 1.46 |

**Table S8. Summary of statistical test to compare the normalized neuronal activity of GPe neurons of cluster3 at saccade onset among conditions during choice task in Figure 3.**

| ***Cluster3*** | | | | | | |
|---|---|---|---|---|---|---|
| parametric bootstrap test (n = 10,000) | *p* | | | | | |
| full model vs. null model | < .001 | | | | | |
| post hoc (pairwise t–test, Bonferroni correction) | Mean (SD) | Mean (SD) | *t* | *p* | 95% CI | effect size |
| **Scene1** | | | | | | |
| (good, contra) vs (bad, contra) | –0.90(1.00) | –1.75(0.85) | 7.12 | <.0001 | [0.61, 1.08] | 1.29 |
| (good, contra) vs (good, ipsi) | –0.90(1.00) | –0.98(1.01) | 0.66 | = 5.11 x 10^–1 | [–0.16, 0.31] | 0.12 |
| (good, contra) vs (bad, ipsi) | –0.90(1.00) | –1.82(0.91) | 7.73 | <.0001 | [0.69, 1.15] | 1.40 |
| (bad, contra) vs (good, ipsi) | –1.75(0.85) | –0.98(1.01) | –6.46 | <.0001 | [–1.00, –0.54] | –1.17 |
| (bad, contra) vs (bad, ipsi) | –1.75(0.85) | –1.82(0.91) | 0.61 | = 5.43 x 10^–1 | [–0.16, 0.31] | 0.11 |
| (good, ipsi) vs (bad, ipsi) | –0.98(1.01) | –1.82(0.91) | 7.07 | <.0001 | [0.61, 1.08] | 1.28 |
| **Scene2** | | | | | | |
| (good, contra) vs (bad, contra) | –1.11(1.02) | –1.81(0.85) | 5.87 | <.0001 | [0.47, 0.93] | 1.06 |
| (good, contra) vs (good, ipsi) | –1.11(1.02) | –1.16(1.05) | 0.41 | = 6.85 x 10^–1 | [–0.19, 0.28] | 0.07 |
| (good, contra) vs (bad, ipsi) | –1.11(1.02) | –1.92(0.87) | 6.76 | <.0001 | [0.57, 1.04] | 1.23 |
| (bad, contra) vs (good, ipsi) | –1.81(0.85) | –1.16(1.05) | –5.46 | <.0001 | [–0.88, –0.42] | –0.99 |
| (bad, contra) vs (bad, ipsi) | –1.81(0.85) | –1.92(0.87) | 0.92 | = 3.59 x 10^–1 | [–0.13, 0.35] | 0.17 |
| (good, ipsi) vs (bad, ipsi) | –1.16(1.05) | –1.92(0.87) | 6.36 | <.0001 | [0.53, 1.00] | 1.16 |
| **Scene3** | | | | | | |
| (good, contra) vs (bad, contra) | –1.06(1.00) | –1.78(0.90) | 6.04 | <.0001 | [0.49, 0.95] | 1.09 |
| (good, contra) vs (good, ipsi) | –1.06(1.00) | –1.18(1.10) | 1.02 | = 3.10 x 10^–1 | [–0.11, 0.36] | 0.18 |
| (good, contra) vs (bad, ipsi) | –1.06(1.00) | –1.69(0.92) | 5.31 | <.0001 | [0.40, 0.87] | 0.96 |
| (bad, contra) vs (good, ipsi) | –1.78(0.90) | –1.18(1.10) | –5.03 | <.0001 | [–0.83, –0.37] | –0.91 |
| (bad, contra) vs (bad, ipsi) | –1.78(0.90) | –1.69(0.92) | –0.73 | = 4.65 x 10^–1 | [–0.32, 0.15] | –0.13 |
| (good, ipsi) vs (bad, ipsi) | –1.18(1.10) | –1.69(0.92) | 4.30 | <.0001 | [0.28, 0.75] | 0.78 |
| **Scene4** | | | | | | |
| (good, contra) vs (bad, contra) | –1.00(0.97) | –1.80(0.95) | 6.76 | <.0001 | [0.57, 1.04] | 1.22 |
| (good, contra) vs (good, ipsi) | –1.00(0.97) | –1.15(1.04) | 1.30 | = 1.94 x 10^–1 | [–0.08, 0.39] | 0.24 |
| (good, contra) vs (bad, ipsi) | –1.00(0.97) | –1.77(0.97) | 6.44 | <.0001 | [0.53, 1.00] | 1.17 |
| (bad, contra) vs (good, ipsi) | –1.80(0.95) | –1.15(1.04) | –5.46 | <.0001 | [–0.88, –0.42] | –0.99 |
| (bad, contra) vs (bad, ipsi) | –1.80(0.95) | –1.77(0.97) | –0.33 | = 7.45 x 10^–1 | [–0.27, 0.20] | –0.06 |
| (good, ipsi) vs (bad, ipsi) | –1.15(1.04) | –1.77(0.97) | 5.14 | <.0001 | [0.38, 0.85] | 0.93 |

**Table S9. Summary of statistical test to compare the normalized neuronal activity of GPe neurons of clusters 1, 2, and 3 among Return, Stay during choice task, and fixation task in Figure 4.**

| ***Cluster1*** | | | | | | |
|---|---|---|---|---|---|---|
| parametric bootstrap test (n = 10,000) | *p* | | | | | |
| full model vs. null model | < .001 | | | | | |
| post hoc (pairwise t–test, Bonferroni correction) | Mean (SD) | Mean (SD) | *t* | *p* | 95% CI | effect size |
| Contra | | | | | | |
| (Return, choice) vs (Stay, choice) | 1.01(1.18) | 0.73(1.00) | 1.55 | = 1.22 x 10^–1 | [–0.08, 0.66] | 0.30 |
| (Return, choice) vs (Good, fixation) | 1.01(1.18) | 1.18(1.13) | –1.21 | = 2.27 x 10^–1 | [–0.73, 0.17] | –0.29 |
| (Return, choice) vs (Bad, fixation) | 1.01(1.18) | 1.09(1.13) | –0.85 | = 3.99 x 10^–1 | [–0.65, 0.26] | –0.20 |
| (Stay, choice) vs (Good, fixation) | 0.73(1.00) | 1.18(1.13) | –2.40 | = 1.72 x 10^–2 | [–1.04, –0.10] | –0.59 |
| (Stay, choice) vs (Bad, fixation) | 0.73(1.00) | 1.09(1.13) | –2.05 | = 4.18 x 10^–2 | [–0.95, 0.02] | –0.51 |
| (Good, fixation) vs (Bad, fixation) | 1.18(1.13) | 1.09(1.13) | 0.33 | = 7.44 x 10^–1 | [–0.59, 0.42] | 0.09 |
| Ipsi | | | | | | |
| (Return, choice) vs (Stay, choice) | 0.41(1.35) | 0.34(1.16) | 0.34 | = 7.38 x 10^–1 | [–0.32, 0.45] | 0.07 |
| (Return, choice) vs (Good, fixation) | 0.41(1.35) | –0.12(1.41) | 1.87 | = 6.31 x 10^–2 | [–0.02, 0.88] | 0.45 |
| (Return, choice) vs (Bad, fixation) | 0.41(1.35) | –0.22(1.30) | 2.30 | = 2.21 x 10^–2 | [0.08, 0.98] | 0.55 |
| (Stay, choice) vs (Good, fixation) | 0.34(1.16) | –0.12(1.41) | 1.48 | = 1.39 x 10^–1 | [–0.12, 0.84] | 0.38 |
| (Stay, choice) vs (Bad, fixation) | 0.34(1.16) | –0.22(1.30) | 1.89 | = 5.95 x 10^–2 | [–0.02, 0.94] | 0.48 |
| (Good, fixation) vs (Bad, fixation) | –0.12(1.41) | –0.22(1.30) | 0.39 | = 6.98 x 10^–1 | [–0.41, 0.60] | 0.10 |
| ***Cluster2*** | | | | | | |
| parametric bootstrap test (n = 10,000) | *p* | | | | | |
| full model vs. null model | < .001 | | | | | |
| post hoc (pairwise t–test, Bonferroni correction) | Mean (SD) | Mean (SD) | *t* | *p* | 95% CI | effect size |
| Contra | | | | | | |
| (Return, choice) vs (Stay, choice) | –1.11(1.08) | –0.92(1.11) | –1.13 | = 2.59 x 10^–1 | [–0.45, 0.12] | –0.18 |
| (Return, choice) vs (Good, fixation) | –1.11(1.08) | 0.35(1.45) | –7.73 | <.0001 | [–1.75, –1.04] | –1.56 |
| (Return, choice) vs (Bad, fixation) | –1.11(1.08) | 0.13(1.12) | –6.50 | <.0001 | [–1.53, –0.82] | –1.31 |
| (Stay, choice) vs (Good, fixation) | –0.92(1.11) | 0.35(1.45) | –6.60 | <.0001 | [–1.60, –0.87] | –1.37 |
| (Stay, choice) vs (Bad, fixation) | –0.92(1.11) | 0.13(1.12) | –5.41 | <.0001 | [–1.38, –0.64] | –1.13 |
| (Good, fixation) vs (Bad, fixation) | 0.35(1.45) | 0.13(1.12) | 1.12 | = 2.65 x 10^–1 | [–0.17, 0.61] | 0.25 |
| Ipsi | | | | | | |
| (Return, choice) vs (Stay, choice) | –1.13(1.22) | –0.99(1.12) | –0.89 | = 3.72 x 10^–1 | [–0.42, 0.16] | 0.14 |

| | | | | | | |
|---|---|---|---|---|---|---|
| (Return, choice) vs (Good, fixation) | –1.13(1.22) | –0.14(1.26) | –5.12 | <.0001 | [–1.28, –0.57] | –1.03 |
| (Return, choice) vs (Bad, fixation) | –1.13(1.22) | –0.08(0.96) | –5.48 | <.0001 | [–1.35, –0.63] | –1.10 |
| (Stay, choice) vs (Good, fixation) | –0.99(1.12) | –0.14(1.26) | –4.21 | <.0001 | [–1.17, –0.42] | –0.89 |
| (Stay, choice) vs (Bad, fixation) | –0.99(1.12) | –0.08(0.96) | –4.55 | <.0001 | [–1.23, –0.49] | –0.96 |
| (Good, fixation) vs (Bad, fixation) | –0.14(1.26) | –0.08(0.96) | 0.32 | = 7.46 x 10^–1 | [–0.45, 0.33] | –0.07 |
| ***Cluster3*** | | | | | | |
| parametric bootstrap test (n = 10,000) | *p* | | | | | |
| full model vs. null model | < .001 | | | | | |
| post hoc<br>(pairwise t–test, Bonferroni correction) | Mean (SD) | Mean (SD) | *t* | *p* | 95% CI | effect size |
| Contra | | | | | | |
| (Return, choice) vs (Stay, choice) | –1.71(0.86) | –1.67(0.84) | –0.34 | = 7.31 x 10^–1 | [–0.36, 0.25] | –0.07 |
| (Return, choice) vs (Good, fixation) | –1.71(0.86) | –0.15(1.11) | –8.62 | <.0001 | [–1.83, –1.15] | –1.85 |
| (Return, choice) vs (Bad, fixation) | –1.71(0.86) | –0.24(1.08) | –8.10 | <.0001 | [–1.74, –1.06] | –1.74 |
| (Stay, choice) vs (Good, fixation) | –1.67(0.84) | –0.15(1.11) | –7.93 | <.0001 | [–1.79, –1.08] | –1.79 |
| (Stay, choice) vs (Bad, fixation) | –1.67(0.84) | –0.24(1.08) | –7.43 | <.0001 | [–1.70, –0.99] | –1.67 |
| (Good, fixation) vs (Bad, fixation) | –0.15(1.11) | –0.24(1.08) | 0.48 | = 6.34 x 10^–1 | [–0.28, 0.46] | 0.11 |
| Ipsi | | | | | | |
| (Return, choice) vs (Stay, choice) | –1.73(0.83) | –1.57(0.86) | –1.39 | = 1.65 x 10^–1 | [–0.10, 0.56] | –0.29 |
| (Return, choice) vs (Good, fixation) | –1.73(0.83) | –0.25(0.93) | –8.13 | <.0001 | [–1.75, –1.07] | –1.75 |
| (Return, choice) vs (Bad, fixation) | –1.73(0.83) | –0.19(0.90) | –8.48 | <.0001 | [–1.81, –1.13] | –1.82 |
| (Stay, choice) vs (Good, fixation) | –1.57(0.86) | –0.25(0.93) | –6.11 | <.0001 | [–1.55, –0.80] | –1.46 |
| (Stay, choice) vs (Bad, fixation) | –1.57(0.86) | –0.19(0.90) | –6.42 | <.0001 | [–1.61, –0.86] | –1.54 |
| (Good, fixation) vs (Bad, fixation) | –0.25(0.93) | –0.19(0.90) | 0.32 | = 7.52 x 10^–1 | [–0.43, 0.31] | 0.07 |

**Table S10. Summary of statistical test to compare the effects of CPP + NBQX injection into GPe during choice task in Figure 5.**

| ***Injection during choice task*** | | | | |
|---|---|---|---|---|
| parametric bootstrap test (n = 10,000) | *p* | | | |
| full model vs. null model | < .0001 | | | |
| post hoc (pairwise t–test, Bonferroni correction) | Mean (ms) (SD) | Mean (ms) (SD) | *z* | *p* |
| CPP+NBQX Contra Good pre vs. post | 193.53 (10.26) | 235.13 (15.33) | 7.77 | <.0001 |
| Saline Contra Good pre vs. post | 190.80 (9.16) | 188.67 (9.89) | –0.42 | = 6.71 x 10^–1 |
| CPP+NBQX Contra Bad pre vs. post | 281.07 (34.01) | 301.87 (51.00) | 3.34 | = 9.00 x 10^–4 |
| Saline Contra Bad pre vs. post | 277.33 (24.25) | 271.60 (22.58) | –0.95 | = 3.43 x 10^–1 |
| CPP+NBQX Ipsi Good pre vs. post | 195.93 (12.03) | 190.60 (11.55) | –1.05 | = 2.93 x 10^–1 |
| Saline Ipsi Good pre vs. post | 196.27 (8.53) | 191.87 (10.93) | –0.87 | = 3.87 x 10^–1 |
| CPP+NBQX Ipsi Bad pre vs. post | 281.00 (33.55) | 266.87 (42.03) | –2.34 | = 1.94 x 10^–2 |
| Saline Ipsi Bad pre vs. post | 279.60 (32.37) | 273.33 (29.78) | –1.03 | = 3.02 x 10^–1 |

**Table S11. Summary of statistical test to compare the effects of CPP + NBQX injection into GPe while monkeys chose actions for Bad object during Choice task in Figure 5.**

| ***Injection into SNr during Choice task*** | | | | |
|---|---|---|---|---|
| Accept Bad object | | | | |
| parametric bootstrap test (n = 10,000) | *p* | | | |
| full model vs. null model | = 1.00 | | | |
| Return for Bad object | | | | |
| parametric bootstrap test (n = 10,000) | *p* | | | |
| full model vs. null model | < .001 | | | |
| post hoc (pairwise t–test, Bonferroni correction) | Mean (%) (SD) | Mean (%) (SD) | *z* | *p* |
| CPP+NBQX Return Contra Bad pre vs. post | 75.89 (14.46) | 43.22 (35.89) | – 12.62 | <.0001 |
| Saline Return Contra Bad pre vs. post | 80.54 (13.98) | 83.76 (7.93) | 1.65 | = 9.87 x 10^–2 |
| CPP+NBQX Return Ipsi Bad pre vs. post | 74.06( 11.99) | 64.75 (21.98) | –3.79 | = 2.00 x 10^–4 |
| Saline Return Ipsi Bad pre vs. post | 77.26 (14.22) | 81.63 (10.62) | 2.09 | = 3.66 x 10^–2 |
| Stay for Bad object | | | | |
| parametric bootstrap test (n = 10,000) | *p* | | | |
| full model vs. null model | < .001 | | | |
| post hoc (pairwise t–test, Bonferroni correction) | Mean (%) (SD) | Mean (%) (S.D.) | *z* | *p* |
| CPP+NBQX Return Contra Bad pre vs. post | 23.94 (14.21) | 56.62 (35.72) | 12.68 | <.0001 |
| Saline Return Contra Bad pre vs. post | 19.46 (13.98) | 15.80 (8.04) | –1.86 | = 6.25 x 10^–2 |
| CPP+NBQX Return Ipsi Bad pre vs. post | 25.58 (12.22) | 29.50 (24.52) | 1.54 | = 1.23 x 10^–1 |
| Saline Return Ipsi Bad pre vs. post | 22.74 (14.22) | 18.37 (10.62) | –2.13 | = 3.28 x 10^–2 |
